# Enzymatically active exudates from *Alteromonas* facilitate *Prochlorococcus* survival in stationary phase

**DOI:** 10.1101/2025.05.28.656624

**Authors:** Zhiying Lu, Sydney Plummer, James Kizziah, Steven J. Biller, J. Jeffrey Morris

## Abstract

The cyanobacterium *Prochlorococcus* has a conspicuously reduced genome causing it to require help from co-existing organisms for survival under a variety of stressful conditions. In this work we demonstrated that the heterotrophic bacterium *Alteromonas macleodii* EZ55 facilitated the survival of *Prochlorococcus* MIT9312 batch co-cultures as they entered stationary phase. We further showed that exudates from both *Alteromonas* and *Prochlorococcus* were responsible for this effect. Unidentified toxic exudates of *Prochlorococcus* lowered the carrying capacity of Pro99 medium for axenic *Prochlorococcus* cells, whereas heat-labile high-molecular weight exudates of *Alteromonas* both removed the effect of *Prochlorococcus* exudates and extended the lifespan of axenic *Prochlorococcus* cultures. *Alteromonas* exudates contained a wide variety of proteins and demonstrated enzymatic activities. Some of these proteins and activities may have been packaged within extracellular membrane vesicles, which we identified within *Alteromonas* exudates and found capable of physically associating with *Prochlorococcus* cells. Many of the functionalities observed in *Alteromonas* exudates (e.g., increasing phosphate availability, degrading hydrogen peroxide) were consistent with leaky Black Queen processes, which are defined as services provided by one organism that benefit the entire community and favor the evolution of interdependencies in microbial communities. Therefore, we discuss the potential ramifications of such processes being packaged into vesicles as opposed to freely diffusing through the extracellular milieu.

## Introduction

In the temperate and tropical oligotrophic ocean, some free-living marine microbes have undergone substantial genome reduction, forcing them to depend on other microbes to help them by providing the products of these lost genes (1). The Black Queen (BQ) Hypothesis is an evolutionary explanation for how genome reduction may be advantageous for free-living microorganisms in well-mixed environments, and may lead members of communities to rely on each other in the absence of kin selection or reciprocation. The BQ Hypothesis states that any biological function that yields products or services that cannot be entirely selfishly retained by the organism performing the function – i.e., the function is *leaky* – provides an opportunity for some lineages in the community to live more efficiently by losing such a function’s genes in order to conserve limited resources (2, 3). Examples of genes encoding leaky functions are those that remove toxins from the extracellular environment (e.g., catalase), produce secreted products (e.g., iron acquisition siderophores), or degrade catabolically desirable polymers in the periplasm or at the cell surface (e.g., alkaline phosphatase, glycolytic enzymes, or peptidases).

The “helper” dynamic between *Prochlorococcus* and *Alteromonas* strains in co-culture experiments is both ecologically relevant and a model for studying BQ evolution (4-8). *Prochlorococcus* is the numerically dominant phytoplankton group throughout the temperate and tropical open ocean gyres, where their small size and efficient metabolism allows them to numerically outcompete larger, more complex phototrophs (9). The high-light ecotypes of *Prochlorococcus* have streamlined genomes lacking enzymes that remove reactive oxygen species (10), leaving them dependent on bacteria like *Alteromonas*, which are found throughout tropical and temperate oceans (11), to mitigate oxidative stress generated by a variety of stressful conditions (12). *Alteromonas* in turn requires *Prochlorococcus* photosynthate to grow in carbon-free culture media. A previous study suggested that removal of hydrogen peroxide was sufficient to explain the growth enhancement effect of co-culturing *Prochlorococcus* with *Alteromonas* (5). However, we were interested to know if *Alteromonas* or *Prochlorococcus* modified the culture environment in other ways that might contribute to, or detract from, *Prochlorococcus* growth. Here, we showed that *Prochlorococcus* cultures also benefited from *Alteromonas*’ help at later stages of batch culture, suggesting the presence of BQ dependencies other than ROS protection. We also showed that most of this helping ability was associated with enzymatically active extracellular products, some of which are likely contained in extracellular membrane vesicles (EVs) that can associate with *Prochlorococcus* cells. These data suggest EVs may serve as a packaging mechanism for preventing the inefficient dilution of BQ functions in dilute oligotrophic planktonic communities.

## Materials and Methods

### Strains and culture conditions

All strains used in this study were taken from those used for a Long-Term Phytoplankton Evolution (LTPE) experiment (13). *Prochlorococcus* strains were streptomycin-resistant derivates of the high light-adapted strain MIT9312 obtained as described previously (4, 5), either before (Ancestor) or after 500 generations of evolution at either 400 ppm or 800 ppm pCO_2_ conditions (i.e., modern day or projected year 2100 conditions (14)). *Alteromonas* strains were derivatives of strain EZ55, originally isolated from a *Prochlorococcus* MIT9215 culture (4). As with our *Prochlorococcus* strains, we used both ancestral and evolved varieties of EZ55 co-evolved with *Prochlorococcus* at the two pCO_2_ treatments and subsequently isolated. *Prochlorococcus* cultures were revived from cultures cryopreserved with 7.5% DMSO in liquid nitrogen vapor, and *Alteromonas* cultures were revived from cultures preserved with 20% glycerol stored at -80° C. Prior to use in experiments, all *Prochlorococcus* cultures were grown in co-culture with *Alteromonas* EZ55 helpers (4) and were acclimated to culture conditions for at least 4 generations prior to data collection.

*Alteromonas* cultures were grown in YTSS medium (15) and *Prochlorococcus* cultures were grown in Pro99 medium (16) or PEv medium (13), both made in an artificial seawater base (ASW) (13). Prior to addition to co-cultures *Alteromonas* strains were pelleted at 2000 g for 2 minutes and washed twice in sterile ASW, then added to cultures at approximately 10^6^ cells ml^-1^. *Alteromonas* was grown at 30° C with 120 rpm shaking. Unless otherwise noted, *Prochlorococcus* and co-cultures were grown in static 13 mL conical bottom acid-washed glass tubes under approximately 75 mmol photons m^-2^ s^-1^ cool white light in a Percival incubator set to 23° C. When medium additions were employed, all solutions were filter sterilized with a 0.2 mm filter. Cell densities of *Prochlorococcus* cultures to standardize inoculations between experiments were determined using a Guava HT1 flow cytometer (Luminex Corporation, Austin, TX) by the distinctive signature of these cells on plots of forward light scatter vs. red fluorescence (Fig. S1A). Day-to-day culture growth was tracked using the *in vivo* chlorophyll *a* module for the Trilogy fluorometer (Turner Designs, San Jose, CA) with a custom 3D-printed adapter designed for conical bottom tubes. Fluorometer measurements and cell counts were linearly related across the range of cells examined in this study (Pearson correlation coefficient 0.835, p = 1.38 x 10^-6^, Fig. S1B).

### Growth tests in conditioned media

We conducted tests using three types of conditioned media: *Prochlorococcus* (Pro CM), *Alteromonas* (EZ55 CM), and *Prochlorococcus* subsequently treated with *Alteromonas* (Pro CM + EZ55). For Pro CM, we produced axenic *Prochlorococcus* by adding streptomycin to a final concentration of 100 μg/mL to low-density (∼10^6^ cells mL^-1^) *Prochlorococcus* cultures. After 48 h exposure to the antibiotic, we confirmed that no *Alteromonas* EZ55 cells survived by transferring 1 mL into sterile YTSS medium and checking for growth after 24 hours. A 0.5 mL aliquot of this axenic *Prochlorococcus* culture was transferred to 12 ml fresh Pro99 media and cultivated for 11 days, after which the cells were removed by filtering the medium using a sterile 0.2 μm PVDF syringe filter (Millipore Sigma, Burlington, MA, USA). EZ55 CM and Pro CM + EZ55 were produced by inoculating washed *Alteromonas* EZ55 cells from YTSS medium at approximately 10^6^ CFU mL^-1^ to sterile Pro99 (EZ55 CM) or to a sub-sample of the Pro CM described above (Pro CM + EZ55). As with Pro CM, these cultures were cultivated for 11 days and were then filtered to remove the cells. To initiate experiments, freshly axenic *Prochlorococcus* (produced as described above) was transferred to replicate 12 mL tubes of each of the 3 conditioned media, and growth was measured by chl-*a* fluorescence every other day.

### *Concentration of* Alteromonas *exudates*

EZ55 was grown in Pro99 media supplemented with 0.1% glucose to sustain growth in the absence of *Prochlorococcus* exudates. We scaled cultures up progressively from 12 mL to 2 L. The 2L culture was grown in a vented bottle with an outlet connected to a filter with 0.22 μm pore size. After removing most of the cells by centrifugation, we produced size-fractionated, concentrated exudates using tangential flow filtration using Sartorius Vivaflow 200 cassettes. The 2L culture supernatant was passed first through a 0.22 μm cassette using a Masterflex L/S peristaltic pump (Cole-Parmer) to remove bacterial cells, then through a 50 kDa module and a 5 kDa module in succession to produce >50 kDa and <50 kDa fractions that were each concentrated approximately 100-fold. A portion of the >50 kDa fraction was placed in boiling water for 5 minutes to denature proteins. When these concentrated extracellular products were added to culture media for growth experiments they were diluted 100-fold, returning them to approximately their original concentration prior to filtration.

### Proteomics

The >50 kDa fraction described above was further concentrated using a 30 kDa centrifugal filter (MilliporeSigma™ Amicon™ Ultra-15, Darmstadt, Germany) to ∼1.5 ml by centrifugation at 7000 g. Then, 13.5 mL sterile milli-Q water was added to the filtrate and was concentrated to ∼1.5 mL again. The above wash step was repeated, and the final ∼1.5 mL sample was transferred to a sterile 2 mL tube for storage at 4° C. We also isolated proteins from whole EZ55 cells from the same cultures used to produce the >50 kDa fraction using a Bacterial Cell Lysis kit (GoldBio). The total protein concentration for each sample was measured using a DC Protein Assay Kit (Bio-Rad, Hercules, CA, USA). The samples were then diluted with 4X Laemmli Sample Buffer (Bio-Rad, Hercules, CA, USA) containing 2-mercaptoethanol (Bio-Rad, Hercules, CA, USA) at the rate of 3 parts sample to 1 part buffer. The diluted sample was heated at 95°C for 5 min, and 20 μL was loaded onto a 4-20% Mini-PROTEAN TGX precast polyacrylamide gel (Bio-Rad, Hercules, CA, USA). Gel electrophoresis was performed in a vertical direction in a Mini-PROTEAN Tetra cell (Bio-Rad, Hercules, CA, USA) at ∼200V for 20-40 min until the blue band in the marker line reached the bottom of the gel. After electrophoresis was complete, the gel was gently removed from the cassette and was rinsed in a shallow staining tray with milli-Q water. The rinsed gel was soaked in fixing solution (40% ethanol, 10% acetic acid) for 15 min with gentle agitation, rinsed with milli-Q water again, and stained with colloidal Coomassie blue for 14 h with gentle agitation at room temperature. The stained gel was destained in three changes of milli-Q water over 3 h with gentle agitation.

For protein identification, the portion of the destained gel containing target bands of interest was cut into 8 slices with equal length (Figure S2), and each slice was digested following the In-Gel Digestion Protocol described by (17). Each digest was analyzed as previously described (18). An aliquot (5 μL) of each digest was loaded onto a Nano cHiPLC 200 μm ID x 0.5 mm ChromXP C_18_ -CL 3-μm 120-Å reverse-phase trap cartridge (Eksigent, Dublin, CA) at 2 μL/min using an Eksigent 415 LC pump and autosampler. After the cartridge was washed for 10 min with 0.1% formic acid in ddH_2_O, the bound peptides were flushed onto a Nano cHiPLC 200-μm ID x 15-cm ChromXP C -CL 3-μm 120-Å reverse-phase column (Eksigent) with a 100-min linear (5 to 50%) acetonitrile gradient in 0.1% formic acid at 1,000 nL/min. The column was then washed with 90% acetonitrile + 0.1% formic acid for 5 min and re-equilibrated with 5% acetonitrile + 0.1% formic acid for 15 min. A Sciex 5600 Triple-TOF mass spectrometer (Sciex, Toronto, Canada) was used to analyze the protein digest. The IonSpray voltage was 2,300 V, and the declustering potential was 80 V. Ion spray and curtain gases were set at 10 and 25 lb/in^2^, respectively. The interface heater temperature was 120°C. Eluted peptides were subjected to a time-of-flight survey scan from m/z 400 to 1250 to determine the top 20 most intense ions for tandem mass spectrometry (MS/MS) analysis. Product ion time-of-flight scans (50 ms) were carried out to obtain the MS/MS spectra of the selected parent ions over the range from m/z 400 to 1,000. The spectra were centroided and deisotoped by Analyst software (v1.7 TF; Sciex). A β-galactosidase trypsin digest was used to establish and confirm the mass accuracy of the mass spectrometer.

The MS/MS data were processed to provide protein identifications using an in-house Protein Pilot 4.5 search engine (Sciex) using the NCBI *Alteromonas* EZ55 protein database and a trypsin digestion parameter and carbamidomethylation for alkylated cysteines as a fixed modification. Proteins of significance were accepted based on the criteria of having at least two peptides detected with a confidence score of >95% using the Paradigm method embedded in the Protein Pilot software. Complete amino acid sequences of predicted proteins were downloaded using the Bio.Entrez package from BioPython (19). Subcellular localization of proteins was predicted using PSORTb v 3.0 (20). KEGG orthology group codes were obtained for proteins using BlastKOALA (21) and were binned into pathways using KEGGREST (22) in R (23). Estimated molecular weights for EZ55 proteins were calculated using the CusaBio molecular weight calculator (https://ww.cusabio.com/m-299.html). Data were statistically analyzed and visualized within R.

### Cryo-electron microscopy (cryo-EM)

Samples of the concentrated and desalted >50kDa fractions were kept in a cool box during transport to the UAB Cryo-EM facility where they were immediately cryogenically frozen in preparation for microscopy as previously described (6). Briefly, 3 μL of each sample were applied to glow-discharged Quantifoil R2/2 200 mesh nickel grids (Electron Microscopy Sciences, Hatfield, PA, USA) and vitrified in liquid ethane using an FEI Vitrobot Mark IV (Thermo Fisher Scientific). The grids were observed on an FEI Tecnai F20 electron microscope (Thermo Fisher Scientific) operated at 200 kV with a typical magnification of ∼34,500 × and 1-3 μm defocus. 111 images were collected under low-dose conditions in SerialEM using a Gatan K3 direct detector. For single particle reconstruction, micrographs were imported to <u>RELION-3</u> (24), and contrast transfer functions were estimated with <u>Gctf</u> (25). Particles were picked with <u>Topaz</u> (26) using a default model (resnet8_u64). A subset of highly symmetric particles were isolated via iterative 2D classification, then separated for 3D reconstruction. Matches to modeled structures were searched in the Protein Data Bank and these structures were then fit to the reconstruction using <u>ChimeraX</u> (27).

### *Exudate interaction with* Prochlorococcus

In order to visualize interactions between exudates and *Prochlorococcus* cells, we covalently labelled the >50 kDa concentrates with an amine-reactive Alexa Fluor 488 5-SDP ester dye (Molecular Probes/ThermoScientific) (28). First, 100 μL of each sample was concentrated with 100 kDa centrifugal ultrafilters (Pall) at 14,000 rpm for 15 minutes and then resuspended with 0.1 M sodium bicarbonate buffer (pH 8.3). Then, 2.5 μL of Alexa Fluor 488 5-SDP ester dye was added and incubated for 1 hour in the dark at room temperature with gentle mixing. The mixture was washed by centrifugation at 14,000 rpm for 15 minutes, removal of the supernatant, and resuspending with 1x PBS three times, with a final resuspension in 100 μL ASW. The labeled and washed exudate was added to 100 μL axenic *Prochlorococcus* culture and incubated for 2h at 28 □ with 120 rpm orbital shaking. Control samples were prepared with *Prochlorococcus* cells only or with *Prochlorococcus* stained with Alexa Fluor 488 5-SDP ester dye but without added exudate concentrate. Cells were analyzed with a Guava HT1 flow cytometer tracking cells by size (forward scatter), chlorophyll (red) fluorescence, and dye (green) fluorescence. Flow cytometry data was visualized using the online Floreada platform (https://floreada.io). Cells were also examined by using a Nikon 80i epifluorescence compound microscope using the GFP and Texas Red filter cubes. Images were collected and processed using Nikon NIS-Elements imaging software.

### Enzyme activities of exudates

We used a variety of enzymatic tests to evaluate the activity of *Alteromonas* exudates. We used the fluorescent conjugates 4-nitrophenyl-α-D-glucopyranoside, 4-nitrophenyl-β-D-glucopyranoside, and 4-nitrophenyl-N-acetyl-β-D-glucopyranoside to measure the activities of the glycolytic enzymes α-glucosidase, β-glucosidase and N-acetyl-glucosaminidase, respectively (29). Phosphatase activity was measured by following the hydrolysis of fluorescent substrate 4-methylumbelliferyl phosphate (30). Enzyme activity for these experiments was expressed as the rate of fluorescent product accumulation over the first 2-5 hours, minus the rate of accumulation in an exudate-free media control. Protease activity was measured after a 30 minute incubation based on Sigma’s non-specific protease activity assay using casein as a substrate (31). Siderophore activity was determined after a 30 minute incubation using chrome azurol S (32). Both fluorescence and optical density traces for these assays were measured using a Biotek Synergy H1 plate reader (Agilent, Santa Clara, CA). Hydroperoxidase activity was assessed by measuring the elimination of a hydrogen peroxide spike after 2 hours, tracking the hydrogen peroxide concentration using acridinium ester chemiluminescence via the injection protocol described previously (5) but modified for use with the Synergy H1. We also attempted to determine the role, if any, of intact EVs in enzyme function by sonicating a subsample of the >50 kDa fraction prior to assays. To accomplish this, the concentrates were treated with a Fisher Scientific sonic dismembrator for 3 cycles of 10 s with 10 s pause between (100W, 30% output efficiency). Both methylumbelliferyl and 4-nitrophenyl conjugate substrates are routinely used to measure membrane permeability due to their inability to cross membranes (33, 34), so an increase in apparent enzymatic activity post-sonication can be interpreted as an indication of membrane disruption.

### Statistical analysis

All statistical analyses were performed in R v. 4.4.1. Most analyses used linear models followed by post hoc extended marginal means testing of pairwise differences between treatment groups using the *emmeans* package (35). Assumptions of linear regression were checked for models by Shapiro-Wilk tests of the normality of residuals and plots of residuals vs. fitted values for homoscedasticity; where these assumptions were violated we used the Box-Cox procedure to find an optimal power transformation (36). Statistical differences between lysate and exudate protein localization counts were determined using Fisher’s exact test implemented in R.

## Results

In our laboratory we routinely maintain *Prochlorococcus* in co-cultures with the helper heterotrophic bacterium *Alteromonas macleodii* EZ55 under low light conditions as seed stocks for experiments. We use two types of media for these cultures: Pro99, a standard medium for *Prochlorococcus* (37), and PEv a low-nutrient medium designed to more closely reflect *Prochlorococcus*’s native oligotrophic habitat (13). Despite the fact that PEv has 1/25 the nutrient concentrations of Pro99, we observed that maximum cell densities in these two media types only differed by approximately 5-fold (Fig. S3), suggesting that stationary phase in Pro99 was not induced by nutrient starvation. We confirmed this by diluting stationary phase Pro99 cultures 2:1 using either Pro99 (Fig. S4) or nutrient free artificial seawater (Fig. S5); in both cases culture growth was nearly identical, with statistically indistinguishable maximum cell densities observed between transfers whether additional nutrients were present or not (linear model, p > 0.05).

To investigate what other factors might have influenced stationary phase cell density, we considered the possibility that the metabolism of dense *Prochlorococcus* cultures may have induced chemical changes in the carbonate system that were toxic to the cells. Like most aquatic phototrophs, *Prochlorococcus* maintains a carbon concentrating mechanism that converts bicarbonate to CO_2_ in the vicinity of Rubisco, consuming protons and raising pH in the process (38). In late exponential stage cultures we observed a density-dependent pH increase of approximately 1 unit from 7.9 to 8.9 (Fig. S6), a 10-fold increase in OH^-^ which would not be substantially alleviated by a 2-fold dilution into fresh media. We also considered the possibility that growth arrest occurred due to carbon limitation, since we grew the cultures in sealed tubes without agitation. However, addition of sterile NaHCO_3_ failed to restore growth, arguing against this possibility (Fig. S7).

As we inspected growth curves of *Prochlorococcus* in Pro99 medium, we observed that many stationary phase cultures experienced pronounced die-offs within a few days of growth cessation (Fig. S4, S5). Because previous studies revealed that axenic *Prochlorococcus* cultures experienced similar mortality events (6, 13, 39), we investigated the role that the co-cultured helper bacterium *Alteromonas* played in stationary phase. Axenic cultures entered stationary phase at significantly lower cell densities than co-cultures with *Alteromonas* (Fig. S8). Moreover, when we diluted stationary phase Pro99 co-cultures 2:1 into ASW as above after the onset of mortality, we still observed restored growth, but not in cultures where *Alteromonas* was killed with antibiotics; these axenic cultures eventually died out completely (Fig. S9). We therefore concluded that *Prochlorococcus* modified the medium in a way that arrested growth and caused cell death prior to nutrient exhaustion, and that *Alteromonas* was able to at least partially ameliorate this condition.

*Alteromonas* and other microbes have been shown to protect dilute *Prochlorococcus* cultures from H_2_O_2_ in their environment, necessitated by the relatively low per-cell ability of *Prochlorococcus* to detoxify this compound (5, 12). However, at higher cell concentrations like those studied here, *Prochlorococcus* was able to bring H_2_O_2_ concentrations below the limit of detection in axenic cultures, indistinguishably from co-cultures with *Alteromonas* (5). We therefore hypothesized that some other product of *Alteromonas* was responsible for extending the growth period and allowing higher maximum cell densities of cultured *Prochlorococcus.* To explore this possibility, we conducted a series of growth experiments to evaluate the effects of both *Prochlorococcus* and *Alteromonas* exudates on the growth of *Prochlorococcus*. We inoculated Pro99 media with either axenic *Prochlorococcus* or one of three *Alteromonas* EZ55 clonal isolates and incubated them under identical conditions for 11 days, then removed all cells by filtration to produce two kinds of conditioned media (Pro-CM and EZ55-CM). We then inoculated a fraction of the Pro-CM with EZ55 cells and incubated them for a further 11 days before removing all cells again by filtration, producing a third type of medium, Pro-CM+EZ55. We then inoculated all three pre-treated media types, as well as unmodified fresh Pro99 as a control, with axenic *Prochlorococcus* cells at a cell density of ∼10^6^ cells mL^-1^ (Fig. 1A). The maximum cell density (Fig. 1B) and area under the growth curve (Fig. S10A) achieved in EZ55-CM was significantly higher than in Pro-CM and usually also higher than fresh Pro99, confirming that bacterial help remained relevant at high cell densities and not just in dilute cultures as previously shown (4, 5). In contrast, exponential growth rates generally did not differ between treatments (Fig. S11A). Also, it was clear that this helping activity was both extracellular (no living *Alteromonas* cells were present in these experiments) and stable for at least a week post-filtration. While the difference was not significant, mean maximum and integrated cell densities in Pro-CM were typically lower than in fresh Pro99 (Fig. 1B, Fig. S10A), consistent with the accumulation of toxic *Prochlorococcus* exudates in denser cultures. In contrast, growth in Pro-CM+EZ55 was not significantly different from growth in EZ55-CM but was usually significantly improved relative to Pro-CM (Fig. 1B), supporting the hypothesis that *Alteromonas* exudates were able to detoxify *Prochlorococcus*’ spent medium.

**Figure 1.**
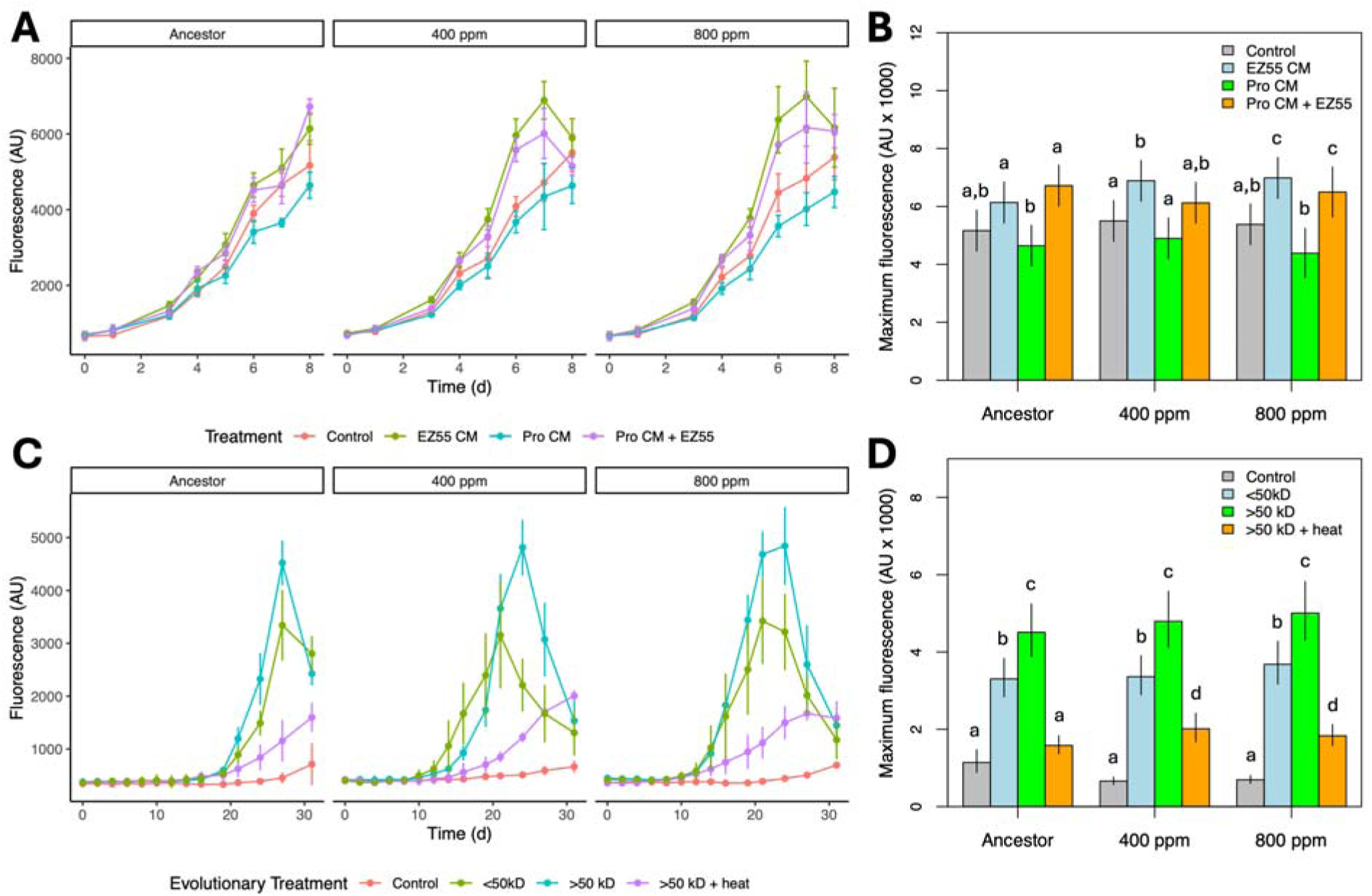
Impact of *Alteromonas* EZ55 exudates on *Prochlorococcus* growth. **(A)** Chlorophyll fluorescence from axenic *Prochlorococcus* cultures grown in Pro99 medium that was either unamended (Control) or else EZ55 (EZ55 CM) or *Prochlorococcus* conditioned either with or without subsequent addition of EZ55 cells (Pro CM and Pro CM + EZ55, respectively). **(B)** Maximum fluorescence achieved by each treatment group and each *Prochlorococcus* evolutionary treatment in Panel A. **(C)** The same *Prochlorococcus* strains grown in Pro99 medium either unamended (Control), with the addition of a concentrated <50KD or >50kD fraction of the cell-free supernatant from an EZ55 culture, or with the >50kD fraction following heat treatment to denature proteins. **(D)** As Panel B, but for the growth curves in Panel C, focusing on the effect of different supernatant size fractions. Letters in panels B and D reflect significance groups from a within-strain pairwise Tukey test. Ancestor, 400 ppm, and 800 ppm reflect *Prochlorococcus* MIT9312 strains before or after 500 generations of evolution under altered pCO_2_ conditions.

We next sought to identify the extracellular *Alteromonas* products contributing to these observations. We size-fractionated and concentrated the contents of all three varieties of EZ55-CM, producing fractions 5-50 kDa (<50 kDa) and >50 kDa. Anticipating that the active components of the higher molecular weight fraction may be proteins or other complex heat-labile molecules, we prepared negative controls by boiling samples of the >50 kDa fraction. We then inoculated dilute (∼1.5 x 10^4^ cells mL^-1^) axenic *Prochlorococcus* cultures in fresh Pro99 medium into which these fractions were added. Under these conditions, unmodified cultures showed no detectable growth for 25 days, consistent with previous studies showing the dependence of dilute *Prochlorococcus* cultures on helpers for survival (4, 5). In contrast, addition of bacterial exudates dramatically enhanced growth (Fig. 1C). The greatest growth enhancement was delivered by the >50 kDa fraction, which was significantly more effective than the smaller molecular weight fraction in raising maximum cell density (Fig. 1D) and also generally more effective for increasing area under the growth curve (Fig. S10B) and exponential growth rate (Fig. S11B). Heat treatment eliminated most of this helping ability, although the boiled samples still allowed greater growth than unmodified Pro99.

Based on these results, we investigated the proteins in the >50 kDa supernatant size fraction and compared them to proteins present in crude cell lysates of the *Alteromonas* strains. Overall, 602 peptides corresponding to predicted *Alteromonas* products were detected (Table S1). A total of 68 proteins were detected in the exudates of all 3 *Alteromonas* isolates, and 144 proteins were detected in the lysates of all isolates (36% and 31% of the total, respectively) (Fig. S12). In contrast, fewer than 10% of proteins were shared between lysates and supernatants (Fig. S13), arguing against the possibility that our supernatant proteins were the result of simple cellular lysis. Surprisingly, fewer than 5% of detected supernatant proteins were predicted to be extracellular (Fig. 2), indicating that most were not canonically secreted proteins. Moreover, 63% of the proteins found in the >50 kDa supernatant fraction were predicted to be smaller than 50 kD, including some as small as 5 kDa (Table S1); this value is only slightly lower than the 71% of proteins <50 kDa found in the crude lysates. Additionally, the average molecular weight of supernatant proteins was only significantly higher than lysate proteins for one *Alteromonas* strain (Fig. S14) and then only by about 20%. The functional categories of proteins were diverse and were not significantly different between lysates and supernatants (Fig. S15).

**Figure 2.**
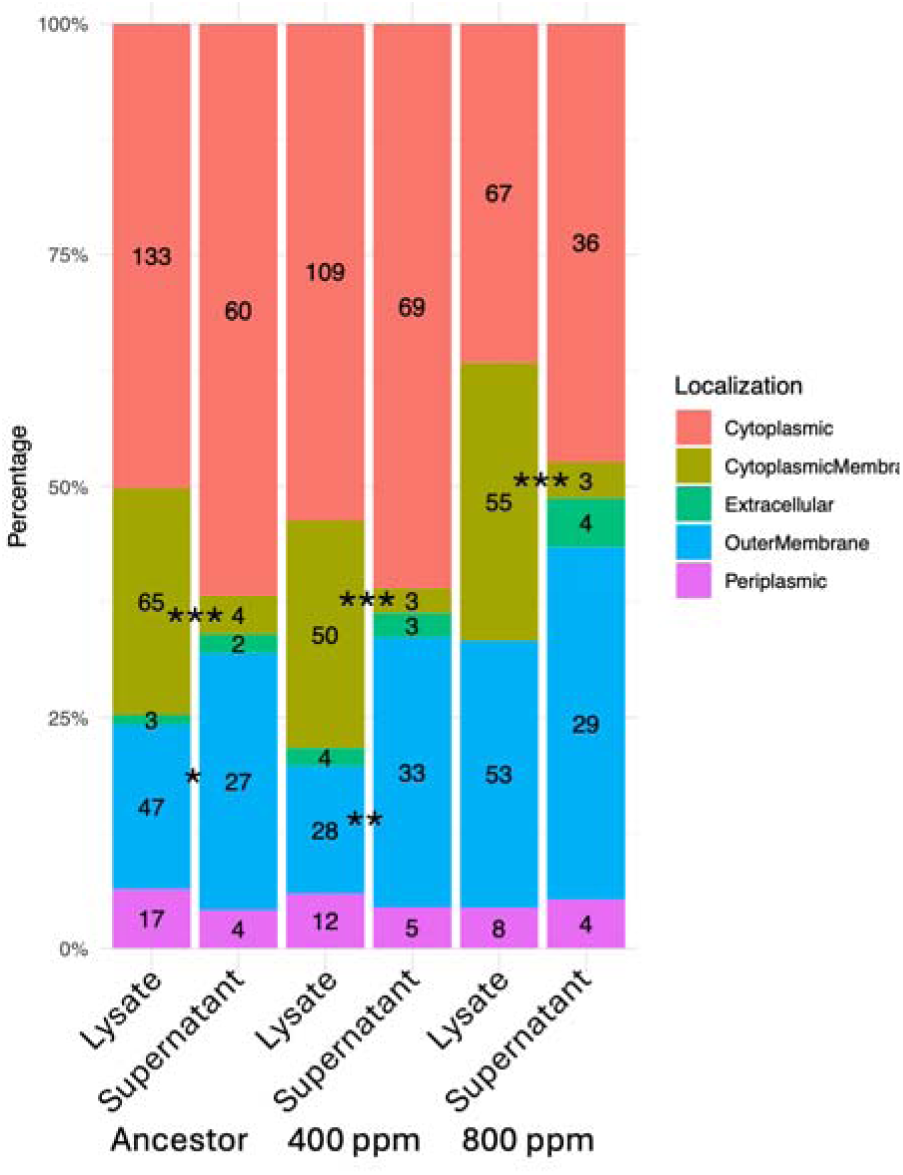
Predicted localization of *Alteromonas* proteins. *Alteromonas* strains were isolated from cultures either before (Ancestor) or after 500 generations of evolution at modern or projected future pCO_2_ conditions (400 ppm and 800 ppm, respectively). Proteins were identified by LC-MS from whole cell lysates or from concentrated cell-free supernatants. Localization of proteins was predicted using psortB. Asterisks indicate the result of Fisher’s Exact Test comparing counts of the indicated proteins between lysate and supernatant for the given *Alteromonas* strain; *, *P* < 0.05; **, *P* < 0.01; ***, *P* < 0.001.

These observations suggested the question of why smaller proteins were retained by the 50 kDa molecular weight filter. We realized that these results might occur if the proteins were packaged inside EVs. EVs are small spherical structures, enclosed by a lipid bilayer, which can contain components from the producing cells. Both gram-positive and gram-negative bacteria can produce EVs (40) and they have been observed previously in both *Prochlorococcus* and *Alteromonas* cultures (28, 41, 42). We observed that proteins predicted to localize to the cytoplasmic membrane were significantly depleted in the supernatant fraction of all *Alteromonas* strains and outer membrane proteins were significantly enriched in 2 of 3 strains (Fisher’s Exact Test, p < 0.05; Fig. 2), consistent with the presence of EVs primarily comprised of the outer membrane of these gram-negative bacteria.

Round or oval EVs between approximately 10 and 100 nm in diameter with clearly visible lipid bilayers were also observed using cryo-EM in all samples (Fig. 3A). Granular particles were observed both within and outside of EVs (Fig. S16). In two cases we were able to use averaged cryo-EM images to identify and reconstruct large symmetrical protein complexes from image data. For the subset of 15nm particles (Fig. 3B), an initial model generated with C2 symmetry displayed higher order icosahedral symmetry, which was applied in subsequent auto-refinement to produce the final reconstruction. The same procedure was applied to the 12nm particles using C4 then D4 symmetry for 3D reconstruction (Fig. 3C). A search of icosahedral structures in the Protein Data Bank identified the 15nm particles as lumazine synthase (PDB: 1NQU), which was fit to the reconstruction with a correlation of 0.97 (Fig. 3D, Fig. S17); the 12nm particles fit a model of apoferritin (the Fe-free structure of bacterioferritin, PDB:1IES) with a correlation of 0.95. The last two steps of riboflavin biosynthesis are catalyzed by lumazine synthase (RibH) and RibE and are found together as a large 60-mer bifunctional icosahedral complex (43) like the one observed here; both of these proteins were detected in the proteomes of all three supernatant samples (Table S1). Similarly, bacterioferritin forms a large cage-like structure comprised of 24 monomers that functions to sequester Fe atoms (44) and was observed in our proteome. While both of these complexes are large enough to be retained by a 50kDa filter, they are also canonically cytoplasmic proteins with no known secretion mechanism. Given the evidence described above that these proteins were not released by simple cell lysis, we suggest that both of these complexes may have been associated with EVs, or have been released through processes similar to those that led to the formation of EVs.

**Figure 3.**
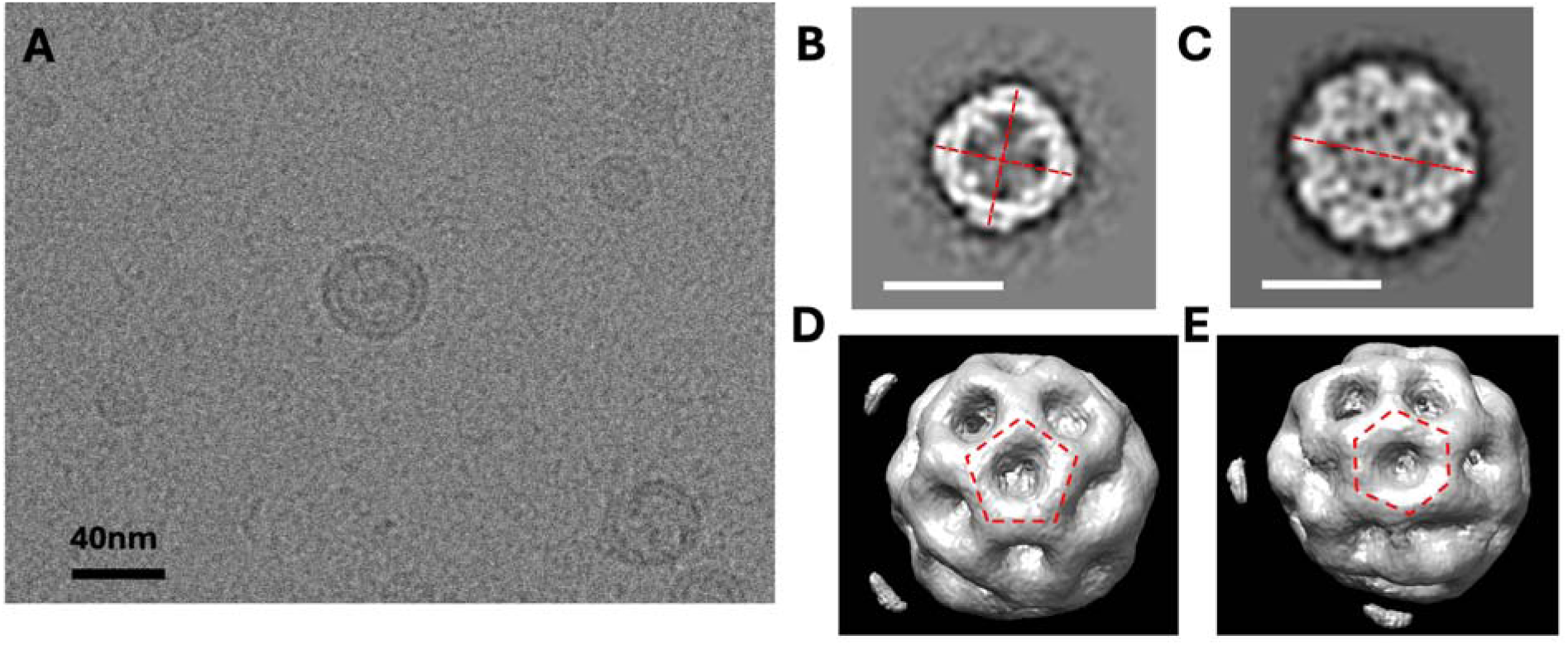
*Alteromonas* exudates contain membrane vesicles and large protein structures. **(A)** Cryo-electron microscopy of concentrated *Alteromonas* exudates revealed abundant bilayer structures resembling membrane vesicles. **(B)** and **(C)** show 2D class average images of 12nm and 15nm particles, respectively, identified by software analysis of cryo-EM images. The dashed lines indicate four- (Panel B) and two-fold (Panel C) axes of rotational symmetry. Panels D and E depict a 3D model obtained by Single Particle Reconstruction of the 15nm particles, showing the characteristic pentagonal **(D)** and hexagonal **(E)** vertices of an icosahedron.

The detection of catalytic enzymes by both proteomics and cryo-EM led us to investigate the enzymatic activity of the >50 kDa supernatant fraction. We detected a-glucosidase and NAGase activity in supernatants collected from the ancestral strain of *Alteromonas* (Fig. S18); b-glucosidase activity was not detected. Protease activity was found in the >50 kDa fraction of all three supernatants but not in the <50kDa fraction; siderophore activity was not significantly different between the size fractions (Fig. S19). Phosphatase and hydroperoxidase (e.g. catalase) activity were detected in all supernatants and were significantly higher in the >50 kDa than the <50 kDa fraction (Fig. 4). We also measured enzyme activity after a sonication treatment was conducted to disrupt the lipid bilayers, reasoning that if the enzymes were packaged in EV lumen, sonication would release them and the observed enzyme activities should increase if the membrane-impermeable substrates no longer had to cross membranes. Significantly increased activities for phosphatase and/or hydroperoxidase were detected post-sonication for all *Alteromonas* strains (Fig. 4).

**Figure 4.**
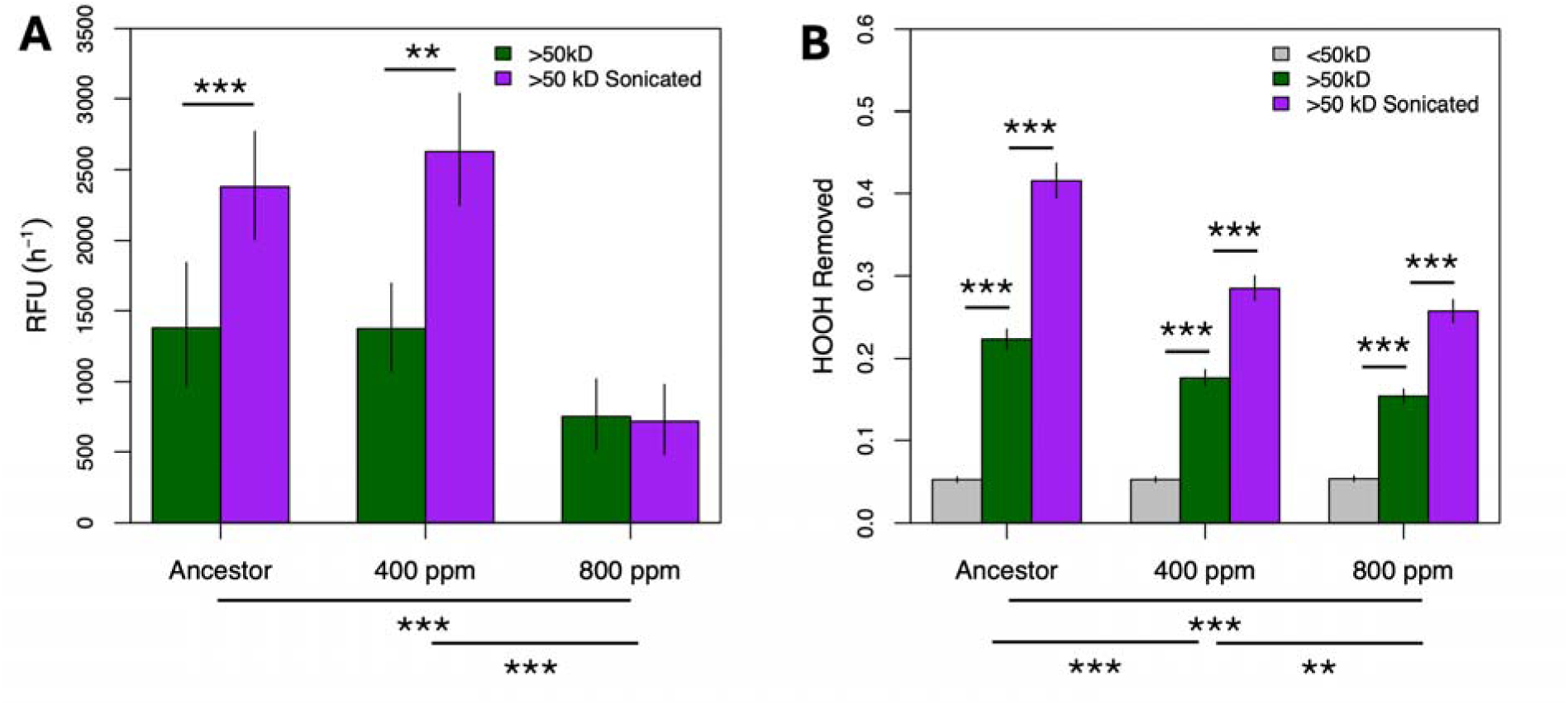
Enzymatic activity in *Alteromonas* exudates. *Alteromonas* strains were isolated from cultures either before (Ancestor) or after 500 generations of evolution at modern or projected future pCO_2_ conditions (400 ppm and 800 ppm, respectively). Cultures were filtered to remove cells and separated into >50kD and <50kD size fractions by tangential flow filtration. Enzyme activity for **(A)** phosphatase and **(B)** hydroperoxidase (e.g., catalase) were assessed for both fractions as well as for the >50kD fraction after a sonication treatment to disrupt vesicles. Phosphatase activity for the <50kD fraction was lower than the blank and was not plotted in Panel A. Asterisks above bars indicate the results of a within-strain Tukey test; asterisks below bars indicate between-strain tests using the average of all treatments. **, *P* < 0.01; ***, *P* < 0.001.

Both intracellular and extracellular vesicles are responsible for transportation and delivery of cargo in eukaryotic systems (45). We reasoned that *Alteromonas* EVs or other extracellular products in our co-culture system might operate in a similar fashion if they tended to physically associate with *Prochlorococcus* cells, delivering leaky functions between cells in a way that could avoid the problems of dilution and degradation in dilute seawater described above. When concentrates from *Alteromonas* >50 kDa supernatant fractions were stained with a green-fluorescing amine-reactive dye and mixed with *Prochlorococcus* cells that emitted red chlorophyll fluorescence and viewed under fluorescence microscopy in multi-color mode, both green and red were found to be associated with some cells (Fig. 5A). Flow cytometry also detected an increase in forward scatter (i.e., size) as well as the appearance of green fluorescence signals for *Prochlorococcus* cells incubated with stained >50 kDa supernatant fractions from all *Alteromonas* strains examined (Fig. 5B-C, Fig. S20). Addition of dye directly to *Prochlorococcus* cells added green signal but did not change the forward scatter profile, suggesting that physical association with exudates rather than transfer of dye to *Prochlorococcus* occurred in these samples. These results indicated the occurrence of direct, physical interactions between *Alteromonas* exudates and *Prochlorococcus* cells when they were incubated together. Moreover, the impact of these associations on the apparent size of the cells is more consistent with aggregation of *Prochlorococcus* with EVs (labeled via covalent attachments between the dye and transmembrane proteins), although we cannot rule out the possibility that other peptides in the exudate also contribute.

**Figure 5.**
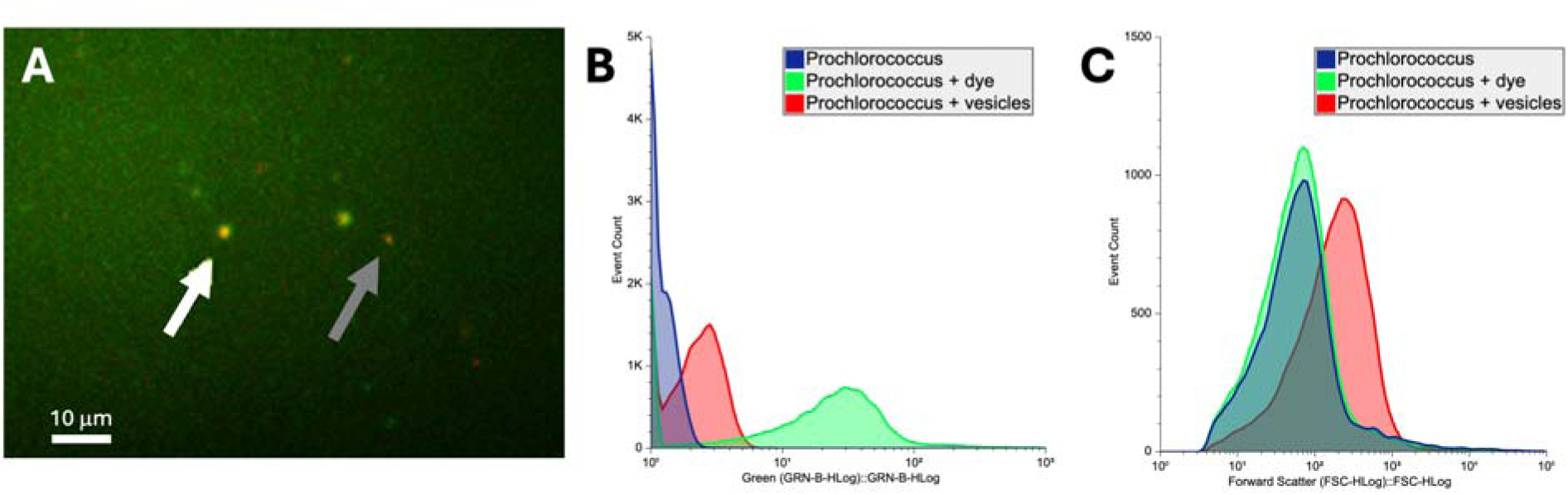
*Alteromonas*-derived exudates physically associate with *Prochlorococcus* cells. **(A)** Exudates labeled with the amine-reactive dye Alexa Fluor 488 were able to stain *Prochlorococcus* cells, suggesting physical association between exudates and cells. The gray arrow indicates an unstained *Prochlorococcus* cell, appearing red from chlorophyll fluorescence; the white arrow indicates a cell also carrying green fluorescence delivered by a stained vesicle. **(B)** Both addition of the dye Alexa 488 as well as dyed *Alteromonas* exudate to *Prochlorococcus* MIT9312 cultures marked the cells with green fluorescence. **(C)** In addition to contributing green fluorescence, addition of dyed exudate (but not Alexa 488 alone) increased the size of MIT9312 cells as measured by forward light scatter, suggesting aggregation of cells and stained vesicles.

Finally, we were interested to know if the composition and activity of exudates was under genetic control in *Alteromonas*. To explore this, we compared *Alteromonas* strains isolated from before (Ancestor) and after 500 generations of evolution at either modern (400 ppm) or projected year 2100 (800 ppm) pCO_2_ conditions (13). The proteins discovered in the supernatant proteome of *Alteromonas* after evolution with *Prochlorococcus* at 800 ppm were notably different than either the ancestral or 400 ppm evolved lineages (Fig. S12). Of the 120 proteins that were not shared between all three strains, 33% were shared between the ancestor and the 400 ppm strain, whereas only 1.7 and 4.2%, respectively, were shared between the 800 ppm strain and the ancestor or the 400 ppm strain. Although the distribution of protein localizations and functional groups were not notably different between the strains (Fig. 2, Fig. S15), we observed significant differences in enzyme activity, with significantly altered a-glucosidase, NAGase, phosphatase, and hydroperoxidase activity in >50 kDa concentrates from the different *Alteromonas* strains (Fig. 4, Figs. S18, S19). Each of these strains was grown in a common garden environment under identical conditions, so these reproducible changes likely reflect genetic differences that arose during evolution (13), suggesting that protein secretion, cellular leakage, and/or vesicle packaging and release were controlled by genetic mechanisms for these organisms.

## Discussion

The dependence of *Prochlorococcus* on other bacteria to help it survive in a variety of conditions has been well established (4-7, 46). Here, we show evidence that *Alteromonas* facilitates greater exploitation of medium resources by *Prochlorococcus*, allowing them to grow to greater cell densities, delay entry into stationary phase, and avoid cell death in batch co-cultures, potentially by degrading autotoxic substances produced in dense *Prochlorococcus* cultures. Moreover, we show that this facilitation is largely accomplished by high molecular weight heat-labile exudates of *Alteromonas* exhibiting diverse enzymatic activities, and these products include, and may be packaged within, EVs. Facilitation of this sort is a central mechanism underlying the BQ Hypothesis that allows reductive genomic evolution and the development of complex exchanges between strains and species in microbial communities (2, 3). Further, this work suggests that EV and protein release are components of this process, at least for the *Prochlorococcus/Alteromonas* interaction. Several previously described BQ functions were observed and/or indicated in cell-free filtrates here, including H_2_O_2_ removal (5, 12), Fe sequestration (47, 48), vitamin biosynthesis (49, 50), and polymer degradation (51-53).

We cannot attribute facilitation exclusively to EVs since fully extracellular proteins were also observed (Fig. 3B, Figs. S16-17). Moreover, many of the putative EV-associated enzymes whose activities we measured (e.g., phosphatase and catalase, Fig. 4), were large enough to have been retained by the 50 kDa filter without being contained in EVs (Table S1), and we observed at least two large extracellular protein complexes by cryo-EM as well (Fig. S16, S17). It is also unclear to what degree these products were actively packaged as opposed to assorted into EVs based on inherent biases in their subcellular location or other biochemical properties, or if instead they were released through uncontrolled processes such as lysis or imperfect cell division. To our knowledge there are no reports of bacterioferritin or lumazine synthase secretion, nor any established mechanisms by which such large complexes could be released from healthy cells. However, the broadly different protein profiles of lysed cells compared to supernatants argued strongly against simply lysis underlying our results, and shifts observed in extracellular proteins following evolution (Fig. 4, Fig. S12) suggested at least some level of regulation acting on protein expression and/or export. Evolutionary changes in bacterial EV characteristics have previously been observed in the Lenski long-term *Escherichia coli* evolution experiment (54) and studies have suggested genetic mechanisms contributing to EV production, secretion, and packaging (55), indicating that EV composition may be a target of selection during bacterial adaptation.

The presence of EVs raises interesting possibilities for the interaction between *Prochlorococcus* and other bacteria specifically, and for BQ interactions more generally. One objection against passive BQ interactions being important in the open ocean is that the average distance between a *Prochlorococcus* cell and the nearest heterotrophic bacterium may be hundreds of cell diameters (56), and if the leaked products are simply excreted into the surrounding environment, their concentrations will exponentially attenuate with distance and will be dramatically diluted by the time they reach potential beneficiaries. Whether the leaked products were metabolites or enzymes, dilution would greatly reduce their impact on their neighbors (56). EVs address this problem in two ways. First, EVs protect secreted materials from degradative environmental factors such as ultraviolet light, suboptimal chemical conditions, and extracellular enzymes, increasing the effective range that secreted material may disseminate while retaining their proper form and function (57, 58). Second, EVs provide a means to keep secreted products close to beneficiary cells once they encounter each other, as they appear to establish stable aggregations with cells (Fig. 5) where fully dissolved proteins may instead remain randomly distributed. Additionally, some fraction of EVs may stay within the diffusive boundary surrounding cells (59), further helping to spatially constrain the shared functions and maintain a locally high relative concentration.

It is also possible that, rather than simply associating with beneficiary cells, EVs instead fuse with the outer membrane, releasing their contents into the periplasm. EV attachment followed by membrane fusion has been proposed as a mechanism for bacterial EVs to deliver compounds to target cells (60-62). Our observations do not preclude the possibility that *Alteromonas* EVs fuse in a similar fashion with *Prochlorococcus* cells. If such events occurred, it would give *Prochlorococcus* at least temporary access to a much wider variety of proteins than its meager genome encodes. While EVs have previously been implicated in genetic transfers between bacteria (63, 64), the interspecies transfer of active enzymes would reflect a form of horizontal information transfer, and suggest a wide expanse of possibilities in understanding metabolism, communication, and cooperation amongst bacteria.

Another intriguing possibility is that EVs may be targeted to associate with particular cells, either of the same species that generated the EVs or with other partners. There is some experimental evidence that this occurs in nature (28, 65) and after laboratory evolutionary differentiation (54) but mechanistic explanations are lacking. While our study was not designed to test this hypothesis, our data nevertheless highlight possible mechanisms for specifying EV delivery. Our proteomic results include numerous proteins predicted to span the outer membrane, including various receptors and transporters (Table S1) that could potentially recognize structures on the outside of other cells or otherwise influence interactions though alterations in surface charge or hydrophobicity. Moreover, we show evidence that *Alteromonas* exudates and possibly also EV packaging changed during the adaptive evolution of *Alteromonas* strains to altered pCO_2_ conditions (Fig. 4, Fig. S18). Previous work also suggested convergent evolution of cell surface features in *Alteromonas* and *Prochlorococcus* strains evolved alongside each other (13). Under the simplest formulation of the BQ Hypothesis, leaky functions yield *public* goods that are freely available to the entire community, but if these functions were packaged, as in EVs, in a manner that could be targeted to specific recipients, these would become *club* goods (i.e., available only to recipients who possessed appropriate credentials in the form of cell surface receptors) and would add another layer of complexity to the evolution of microbial metabolic markets. Future work should investigate the mechanism underlying the physical association between EVs and cells in this system.

While our results show that enzymatically active *Alteromonas* exudates alter Pro99 media in a way that improves *Prochlorococcus* growth and survival under conditions of nutrient exhaustion, it remains unclear what activity is responsible for this facilitation. Our experiments with conditioned media strongly suggest that at least one compound secreted by *Prochlorococcus* caused growth cessation and cell death in Pro99 cultures, and that this compound was deactivated or degraded by *Alteromonas* exudates (Fig. 1, Fig. S9). Several other studies of spent *Prochlorococcus* culture media have revealed a wide variety of metabolites (66, 67) as well as high exudation rates in stationary phase (68), but none were obviously toxic. *Prochlorococcus* secretes several organic acids, but they were not sufficiently concentrated to overcome the alkalinizing effect of the photosynthetic carbon concentrating mechanism (Fig. S6) and were therefore unlikely to be able to cause intracellular acid stress. Another possibility is that one or more extracellular compounds signal *Prochlorococcus* to undergo programmed cell death (PCD). PCD-like activity has been observed in natural populations of *Prochlorococcus* and *Synechococcus* following a defined diel pattern (69), and has also been seen in numerous other species of cyanobacteria and algae (70-77). If *Prochlorococcus* does undergo PCD after reaching a certain cell concentration, this suggests two important questions for further investigation: what benefit does it gain from limiting its own growth while leaving unexploited nutrients in the medium, and what benefit does *Alteromonas* gain from staving this process off for a few more generations?

A notable limitation of the present study is that most of the observations were collected under conditions that do not closely resemble those in the oligotrophic gyres where *Prochlorococcus* is normally found. The nutrient concentrations in Pro99 are orders of magnitude higher than those in the open ocean, and cell densities in these experiments ranged from approximately 10^7^ to 10^8^ cells mL^-1^, whereas maximum cell densities in the ocean do not generally exceed 10^5^ cells mL^-1^ (78). Moreover, *Prochlorococcus* densities in the ocean are likely kept below their carrying capacity by grazing mortality (79), leading to continuous nutrient-limited growth rather than the boom-and-bust dynamics of batch culture. Therefore, while expanding these results into an ecological context is challenging, future work investigating production and activity of *Alteromonas* extracellular products including EVs and their impact on *Prochlorococcus* under continuous culture conditions may better clarify their role in native marine processes.

In conclusion, *Alteromonas* secretes products that reduce *Prochlorococcus* mortality in stationary phase and allow several more generations of growth in Pro99 medium, and some of these products may be associated with EVs. The possibility that the proteins responsible for the BQ interaction between *Prochlorococcus* and *Alteromonas* are packaged in EVs opens up new possibilities for the evolution of these dynamics, including the possibility that the benefits from leaky functions may be targeted to specific community members instead of the community as a whole, making such benefits club goods instead of public goods in economic parlance. Future laboratory experiments should investigate this possibility in this and other systems, and modeling efforts should explore the range of impacts on community evolution and the origin of mutualisms for such targetable leaky functions.

## Acknowledgements

We are grateful to Erik Zinser for providing the ancestral strains used in this study; Zane Forbus for laboratory assistance; Terje Dokland and the UAB Cryoelectron Microscopy Facility; Steven Barnes and Landon Wilson at the UAB Targeted Metabolomics and Proteomics Laboratory for mass spectrometry services; and Luis Zaman for creating the 3D printed tube adapter for the Trilogy fluorometer.

---

**Supplemental Figure 1.**
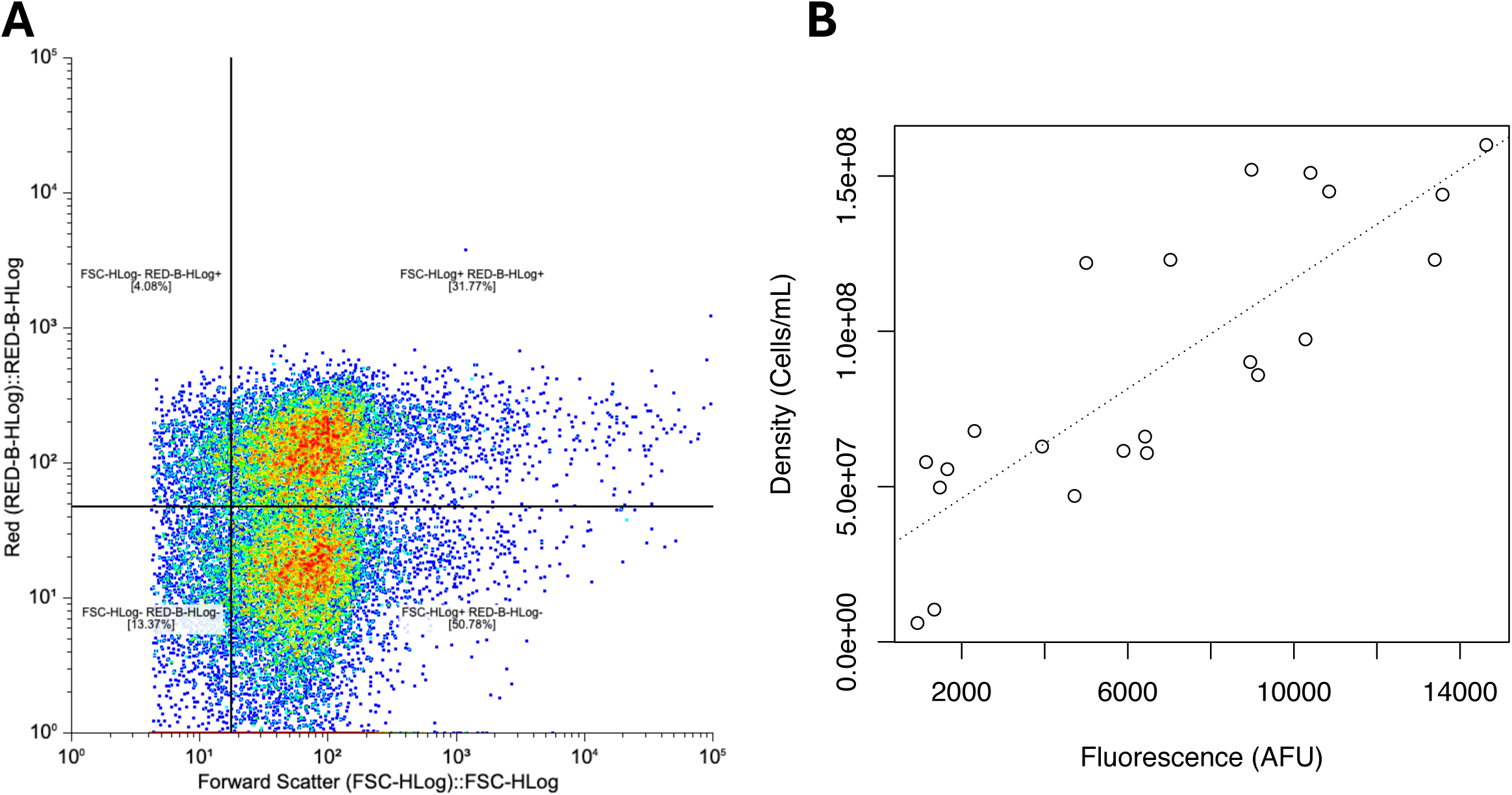
Estimation of *Prochlorococcus* cell density. **(A)** *Prochlorococcus* cultures were enumerated using flow cytometry with gates set on forward scatter and red fluorescence as shown. Events in the upper right quadrant were selected as *Prochlorococcus*. **(B)** Some cultures were tracked by bulk chlorophyll fluorescence, which was a robust linear predictor of cell density within the ranges and conditions employed in these experiments. Dashed line, regression of density on fluorescence for cultures simultaneously assessed for each during growth under identical conditions.

**Supplemental Figure 2.**
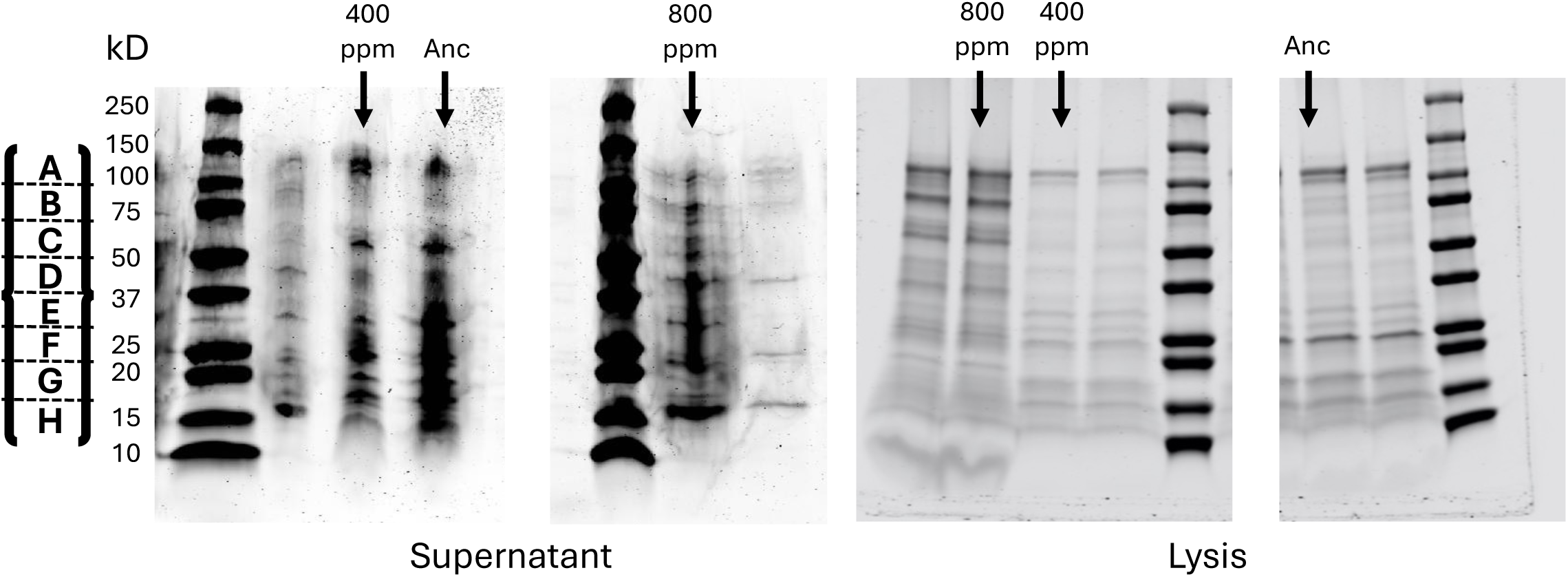
Extracted proteins of *Alteromonas* strains before and after evolution. *Alteromonas* strains were isolated from cultures either before (Ancestor) or after 500 generations of evolution at modern or projected future pCO_2_ conditions (400 ppm and 800 ppm, respectively). Supernatant proteins were obtained by tangential flow filtration with a 50kD filter; lysis proteins were obtained using a cell lysis kit without further concentration. The A-H scheme on the left hand side of the figure shows the approximate locations of the gel slices that were used for LC-MS peptide identification.

**Supplemental Figure 3.**
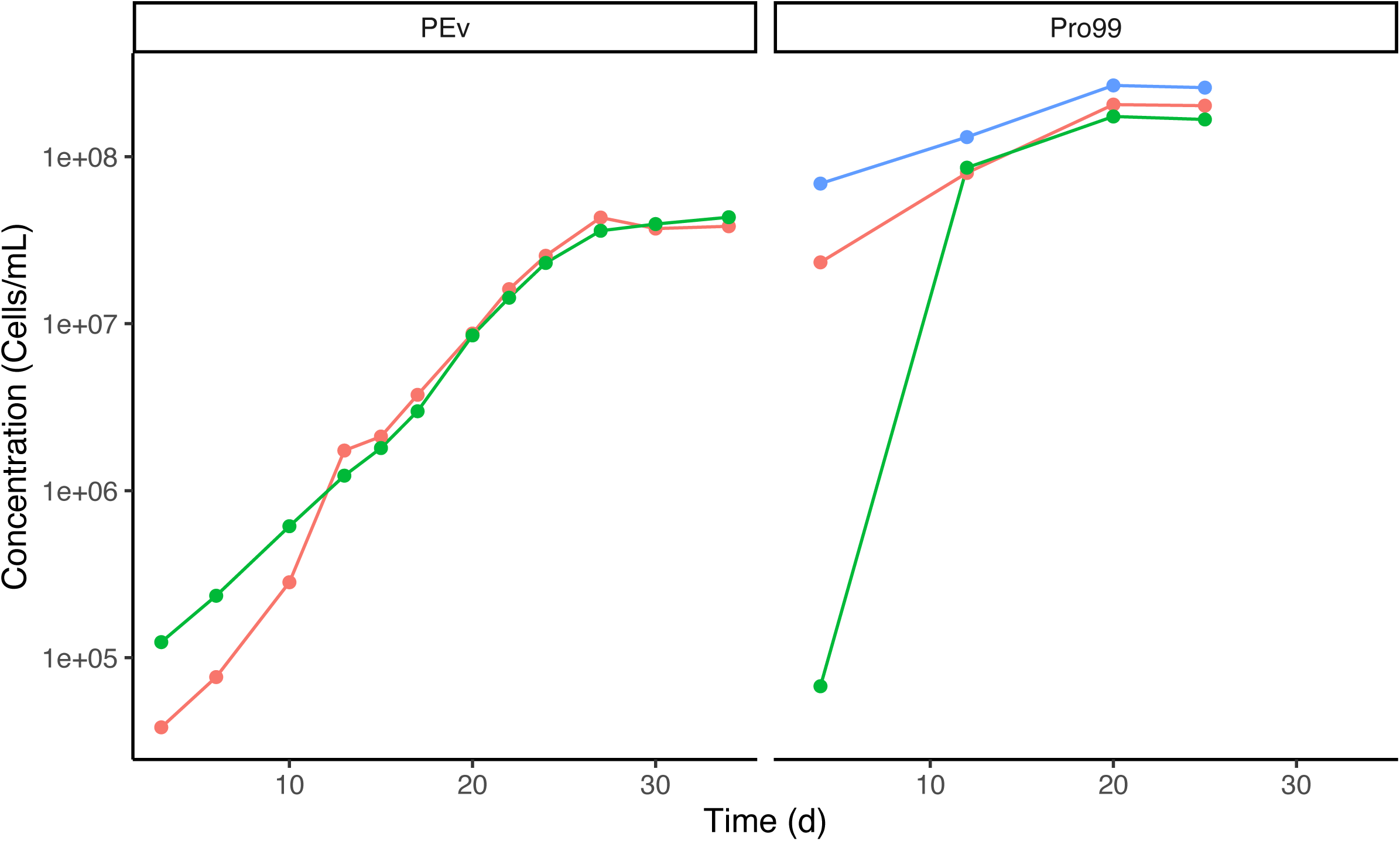
*Prochlorococcus* growth in different media. *Prochlorococcus* growth was tracked by flow cytometry for approximately one month during growth at low (∼30 mmol photons m^-2^ s^-1^) light in either PEv or Pro99 media. Different color lines represent different replicate cultures.

**Supplemental Figure 4.**
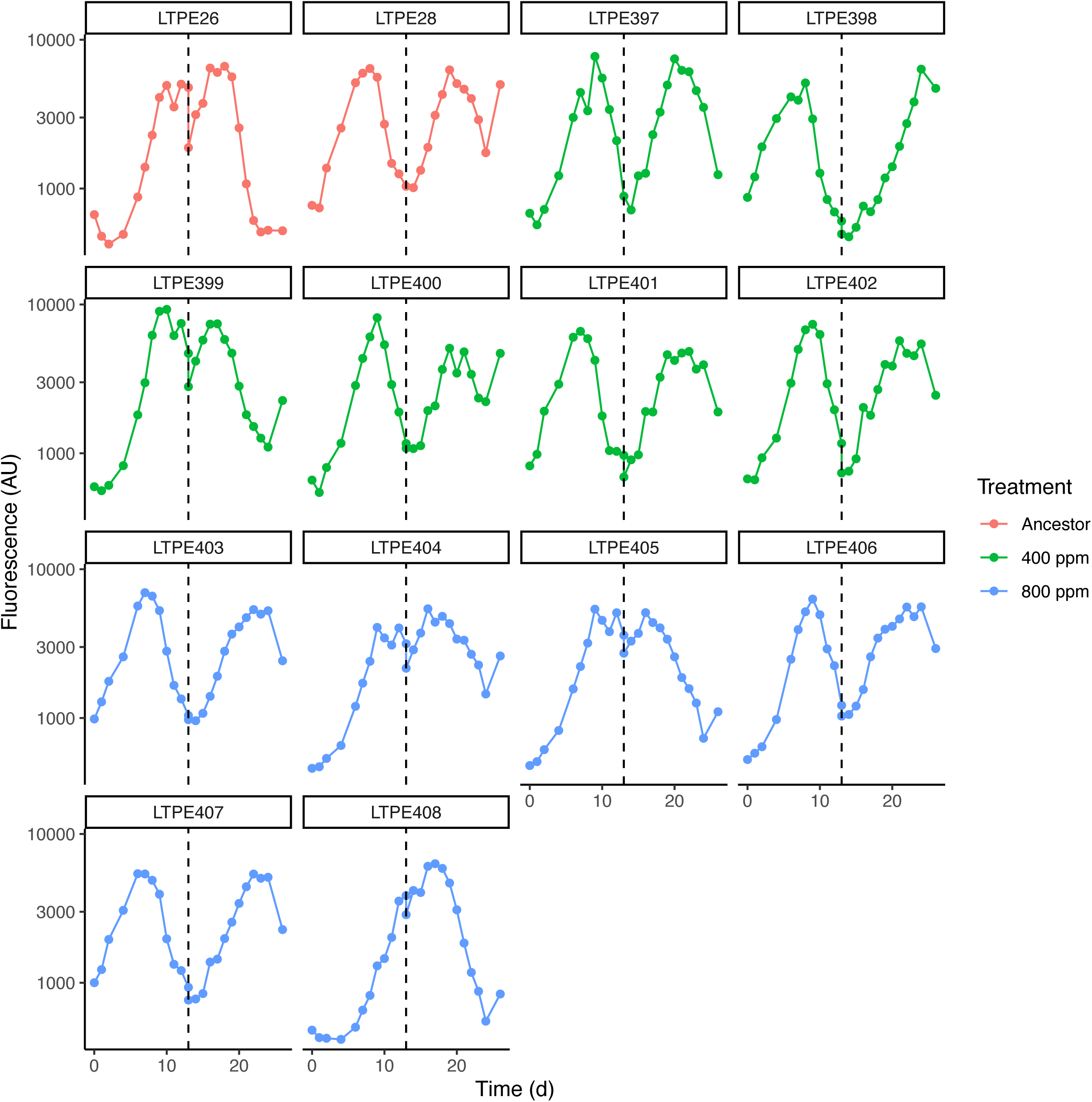
Growth of *Prochlorococcus* after dilution into Pro99 medium. *Prochlorococcus* was allowed to grow to stationary phase and then diluted 2-fold into fresh Pro99 medium at the time point indicated by the dashed line. Each panel represents a replicate culture from the LTPE experiment with evolutionary treatment indicated by color.

**Supplemental Figure 5.**
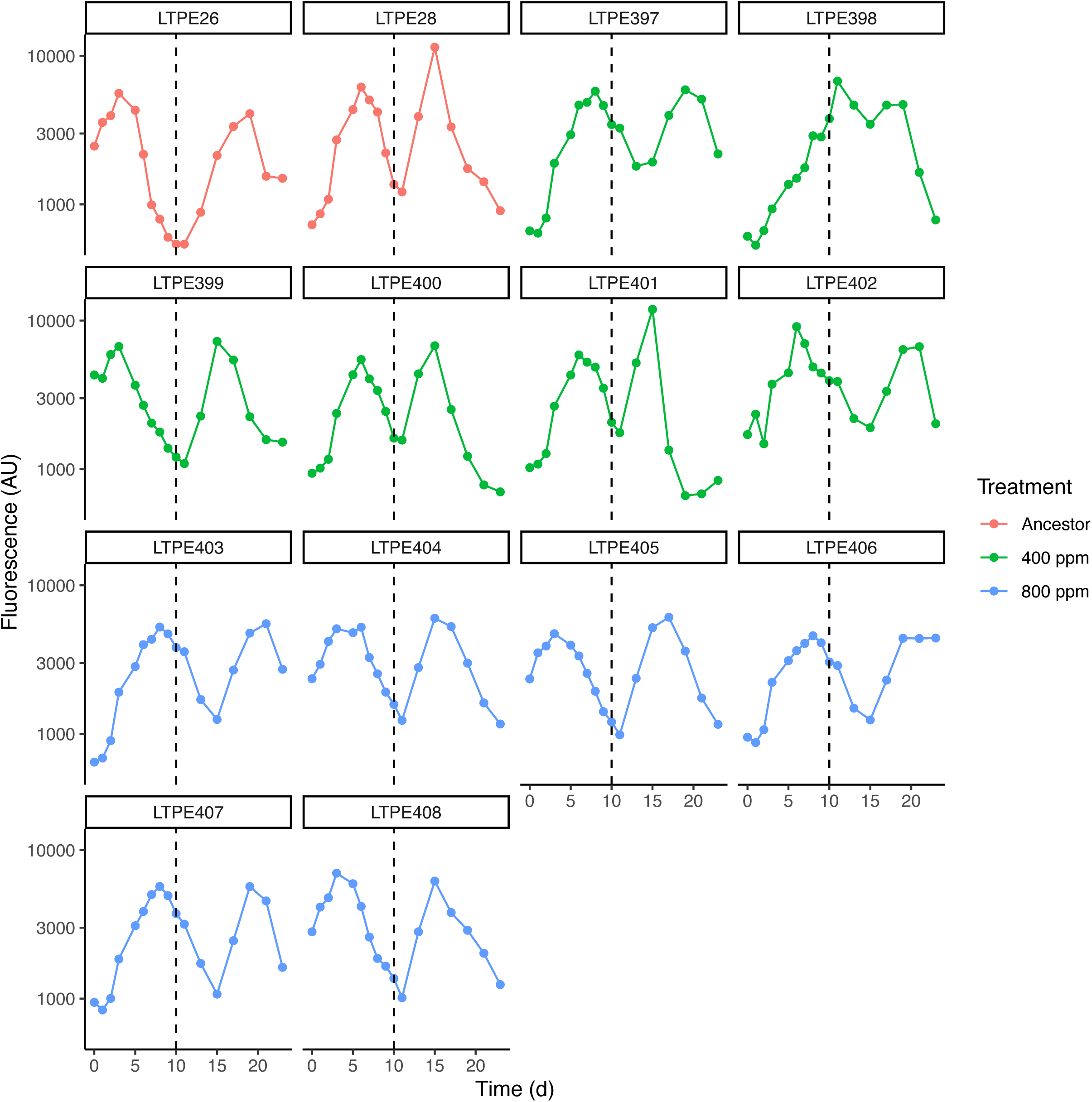
Growth of *Prochlorococcus* after dilution into nutrient-free ASW. *Prochlorococcus* was allowed to grow to stationary phase and then diluted 2-fold into ASW without added nutrients at the time point indicated by the dashed line. Each panel represents a replicate culture from the LTPE experiment with evolutionary treatment indicated by color.

**Supplemental Figure 6.**
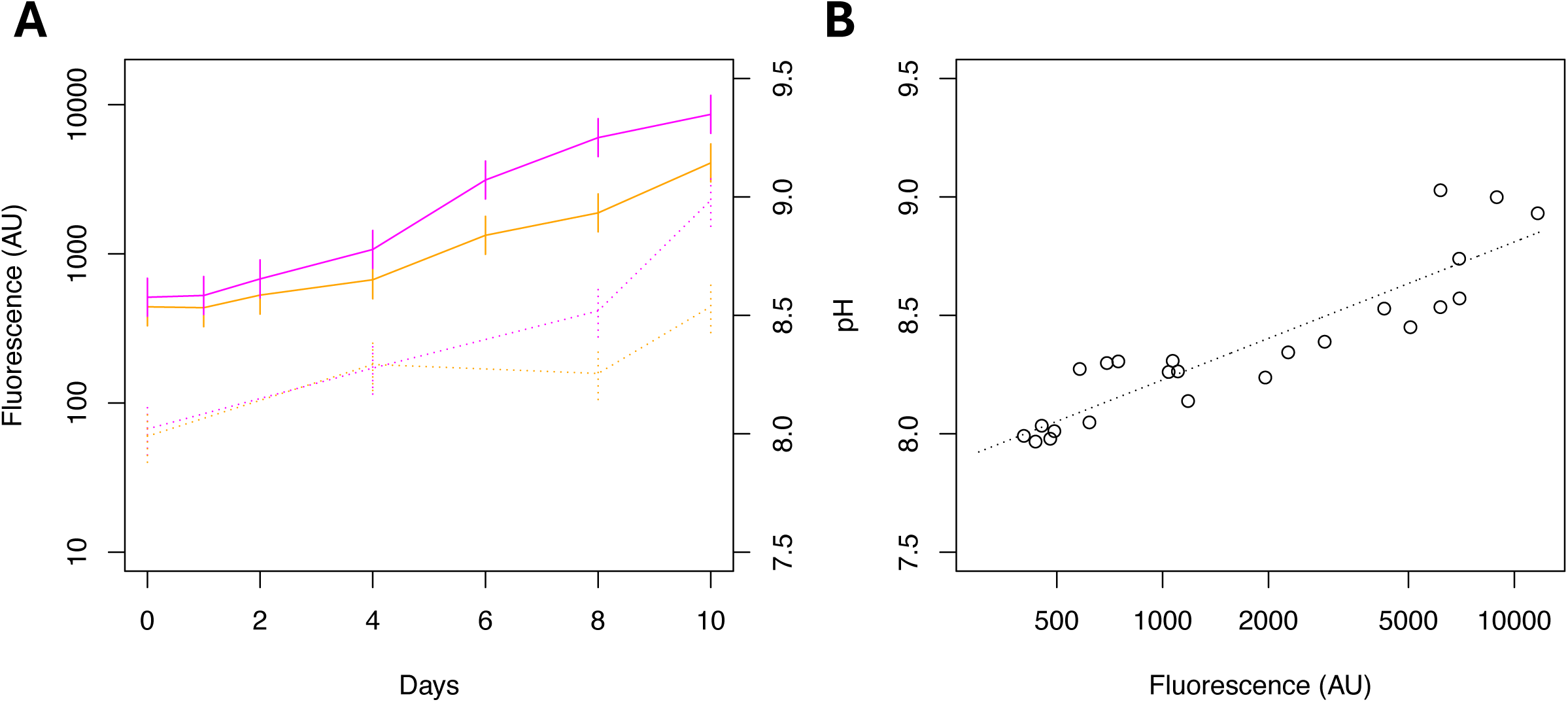
Impact of *Prochlorococcus* growth on medium pH. **(A)** Growth of *Prochlorococcus* (solid lines, left axis) and medium pH (dotted lines, right axis) were measured over the course of a batch culture for three replicates each of LTPE31 (ancestor) and LTPE405 (evolved at 800 ppm pCO_2_). **(B)** Log cell density and medium pH were linearly related over the ranges observed in these experiments (slope of regression of pH on log fluorescence 0.253, r^2^ = 0.81, p = 1.31 x 10^-9^).

**Supplemental Figure 7.**
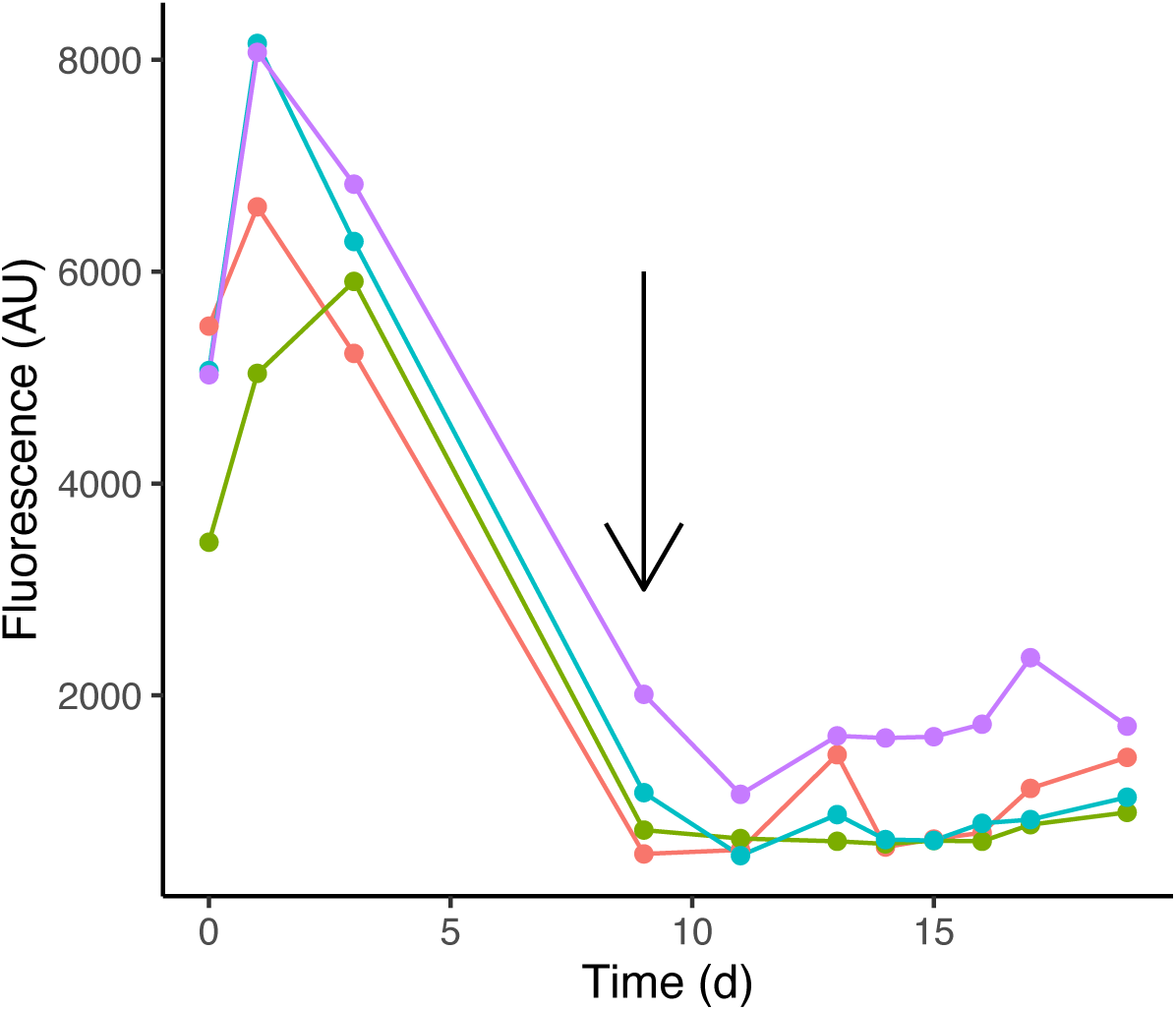
*Prochlorococcus* is not carbon-limited at stationary phase in Pro99. *Prochlorococcus* was allowed to grow to stationary phase in Pro99; when fluorescence values began to drop, NaHCO_3_ was added (at the time point indicated by the arrow). Different color lines represent different replicate cultures from the LTPE experiment.

**Supplemental Figure 8.**
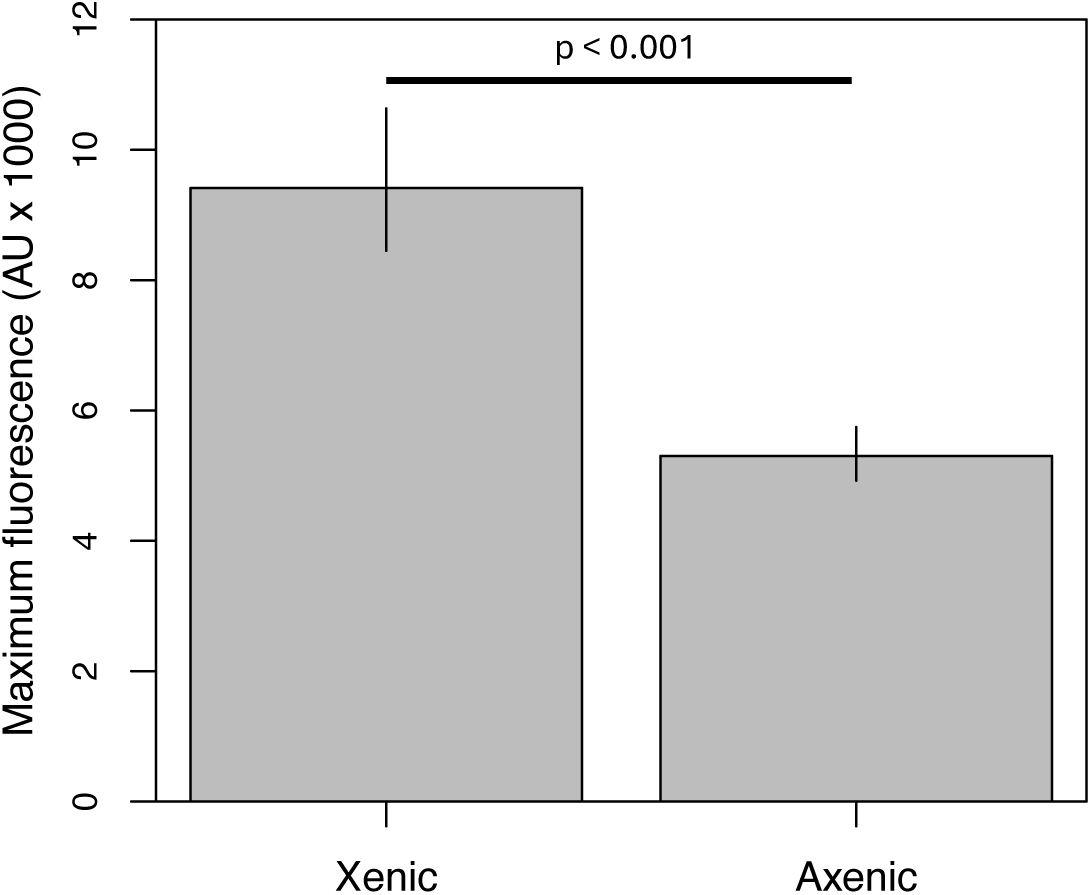
Impact of *Alteromonas* on *Prochlorococcus* carrying capacity. *Prochlorococcus* cultures either with (Xenic) or without (Axenic) *Alteromonas* was allowed to grow into stationary phase to observe their maximum achieved fluorescence. Axenic cultures entered stationary phase at a significantly lower cell density than co-cultures.

**Supplemental Figure 9.**
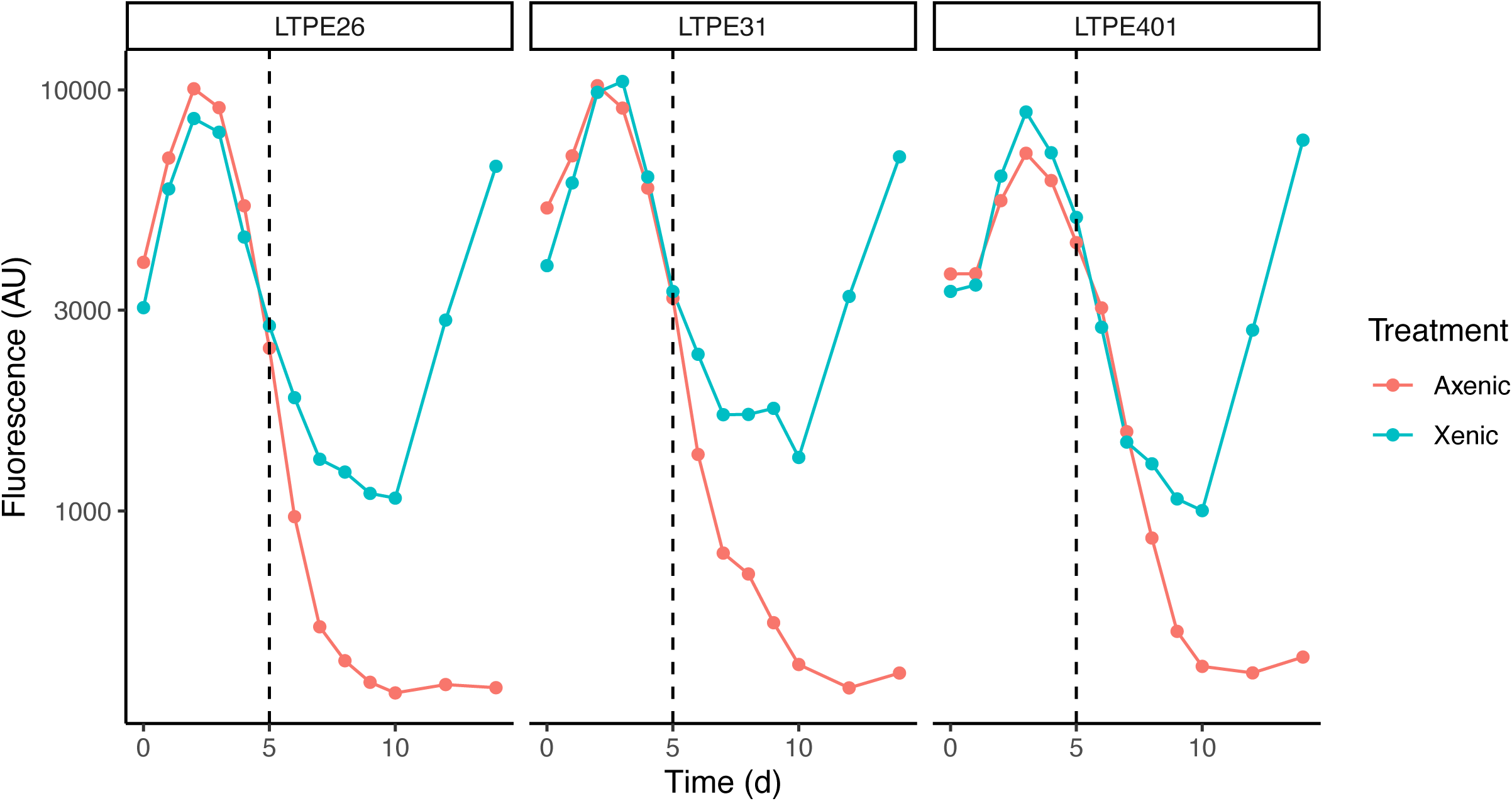
Impact of *Alteromonas* on *Prochlorococcus* survival after transfer. *Prochlorococcus* was grown in co-culture with *Alteromonas* into late stationary phase where fluorescence values began to drop from their peak. Cultures were then diluted 2-fold into ASW without added nutrients at the time point indicated by the dashed line, either with (Axenic) or without (Xenic) the addition of streptomycin to kill *Alteromonas*. Each panel represents a replicate culture from the LTPE experiment; LTPE26 and LTPE31 were ancestral cultures and LTPE401 was from a culture evolved at 400 ppm pCO_2_.

**Supplemental Figure 10.**
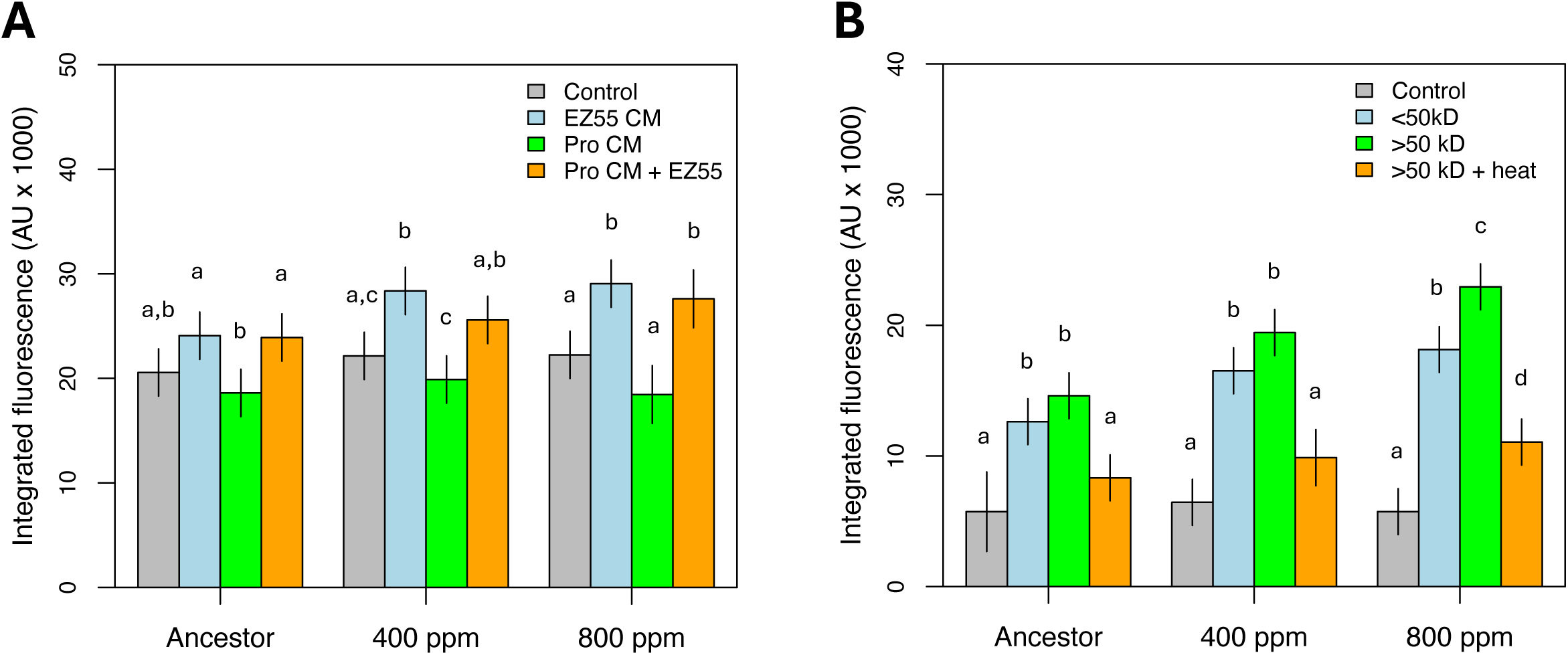
*Prochlorococcus* integrated abundance following treatment with *Alteromonas* exudates. Panels **(A)** and **(B)** reflect the area under the growth curve of the *Prochlorococcus* cultures depicted in Main Text Figures 1A and 1B, respectively. Letters reflect significance groups from a within-strain pairwise Tukey test. Ancestor, 400 ppm, and 800 ppm reflect *Prochlorococcus* MIT9312 strains before or after 500 generations of evolution under altered pCO_2_ conditions.

**Supplemental Figure 11.**
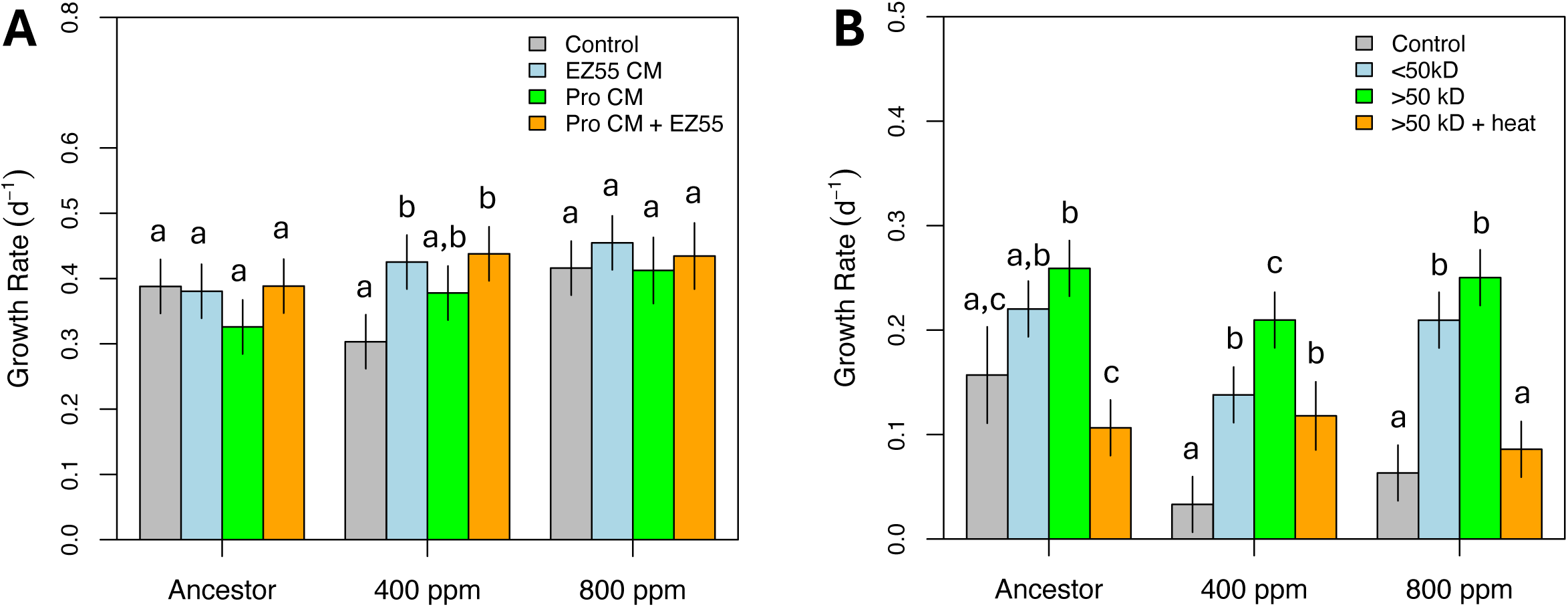
*Prochlorococcus* growth rates following treatment with *Alteromonas* exudates. Panels **(A)** and **(B)** reflect the exponential growth rates of the *Prochlorococcus* cultures depicted in Main Text Figures 1A and 1B, respectively. Letters reflect significance groups from a within-strain pairwise Tukey test. Ancestor, 400 ppm, and 800 ppm reflect *Prochlorococcus* MIT9312 strains before or after 500 generations of evolution under altered pCO_2_ conditions.

**Supplemental Figure 12.**
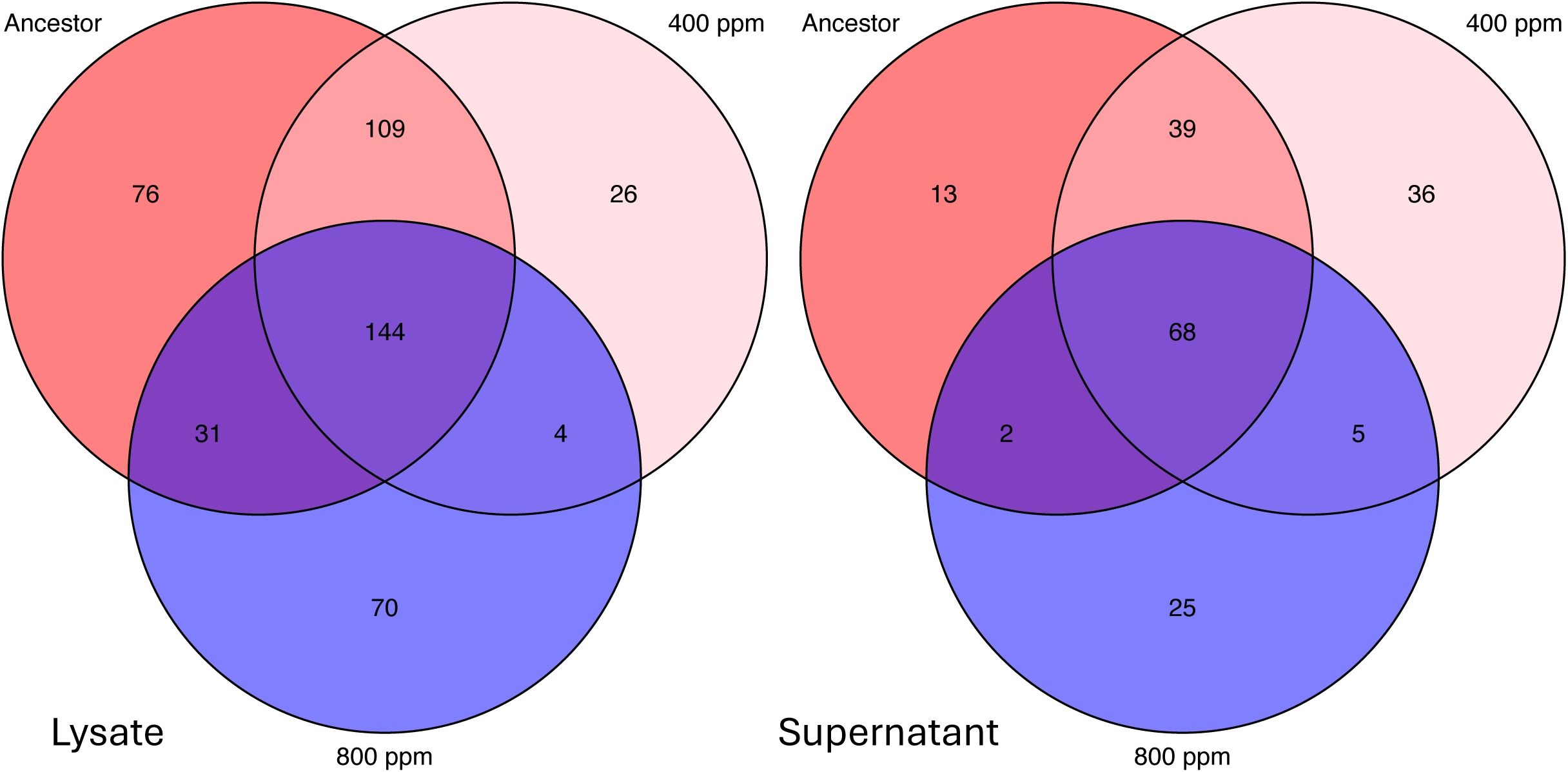
Proteome of *Alteromonas* strains before and after evolution. *Alteromonas* strains were isolated from cultures either before (Ancestor) or after 500 generations of evolution at modern or projected future pCO_2_ conditions (400 ppm and 800 ppm, respectively). Proteins were identified by LC-MS from whole cell lysates or from concentrated cell-free supernatants. These Venn diagrams show the number of shared proteins between strains for each preparation treatment.

**Supplemental Figure 13.**
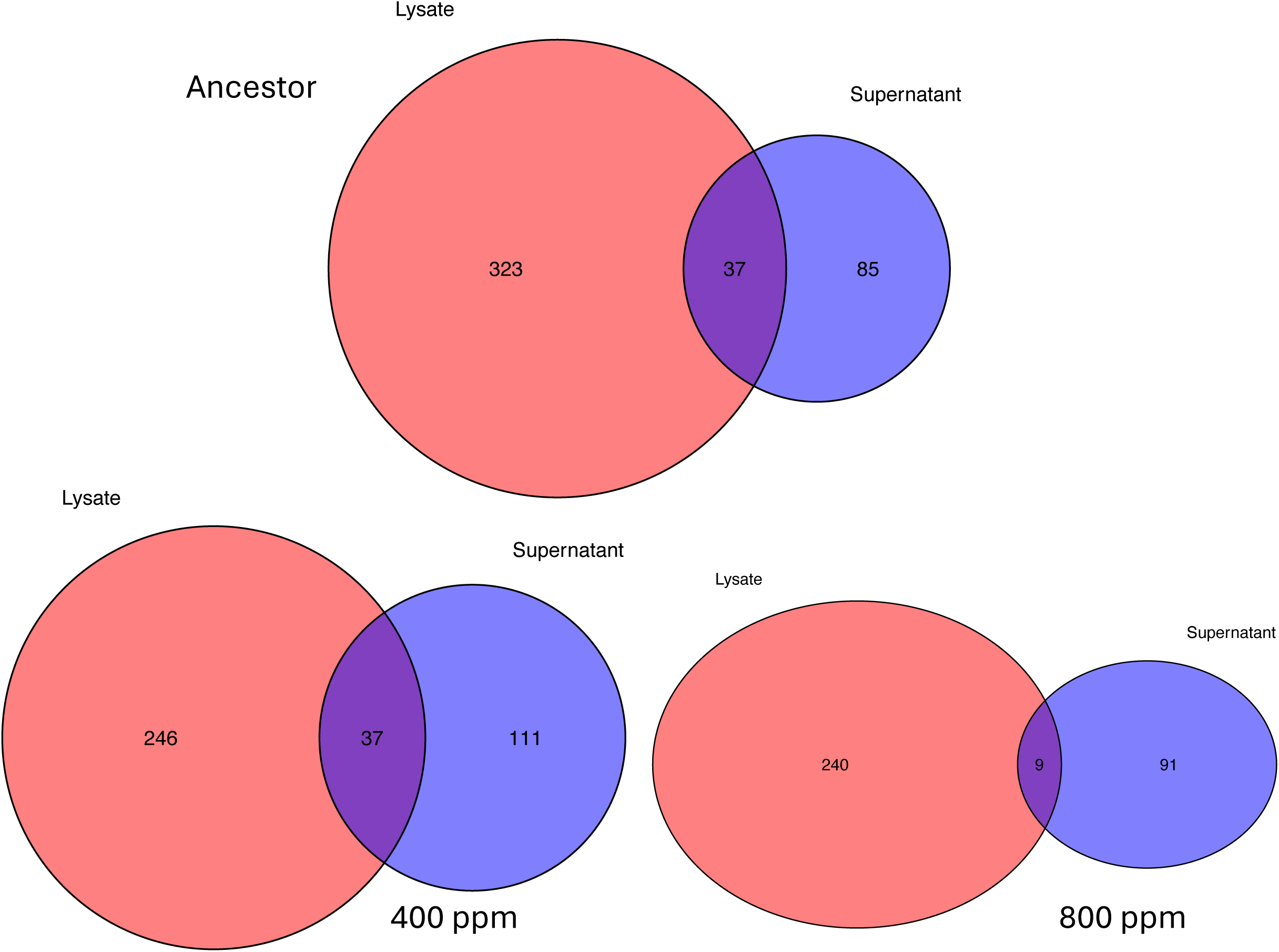
Overlap of *Alteromonas* intracellular and extracellular proteomes. *Alteromonas* strains were isolated from cultures either before (Ancestor) or after 500 generations of evolution at modern or projected future pCO_2_ conditions (400 ppm and 800 ppm, respectively). Proteins were identified by LC-MS from whole cell lysates or from concentrated cell-free supernatants. These Venn diagrams show the shared proteins between intracellular (lysate) and extracellular (supernatant) treatment groups for each type of strain.

**Supplemental Figure 14.**
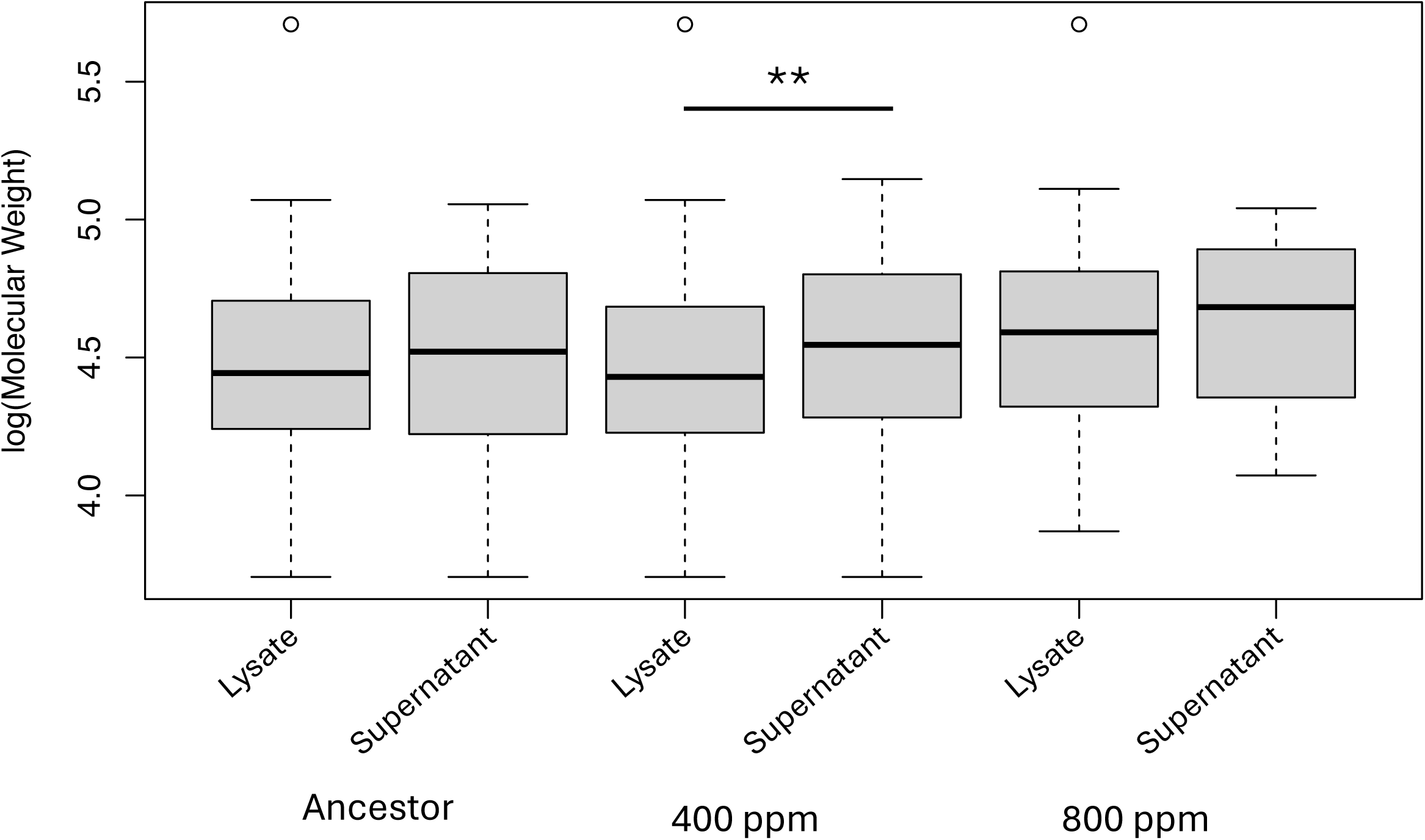
Molecular weights of proteins from *Alteromonas* exudates. *Alteromonas* strains were isolated from cultures either before (Ancestor) or after 500 generations of evolution at modern or projected future pCO_2_ conditions (400 ppm and 800 ppm, respectively). Proteins were identified by LC-MS from whole cell lysates or from concentrated cell-free supernatants and molecular weights were predicted using corresponding sequences from the *Alteromonas* EZ55 genome. Asterisks indicate *P* < 0.01 from of a post-hoc t-test comparing the mean molecular weight of lysate and supernatant proteins for each EZ55 strain category.

**Supplemental Figure 15.**
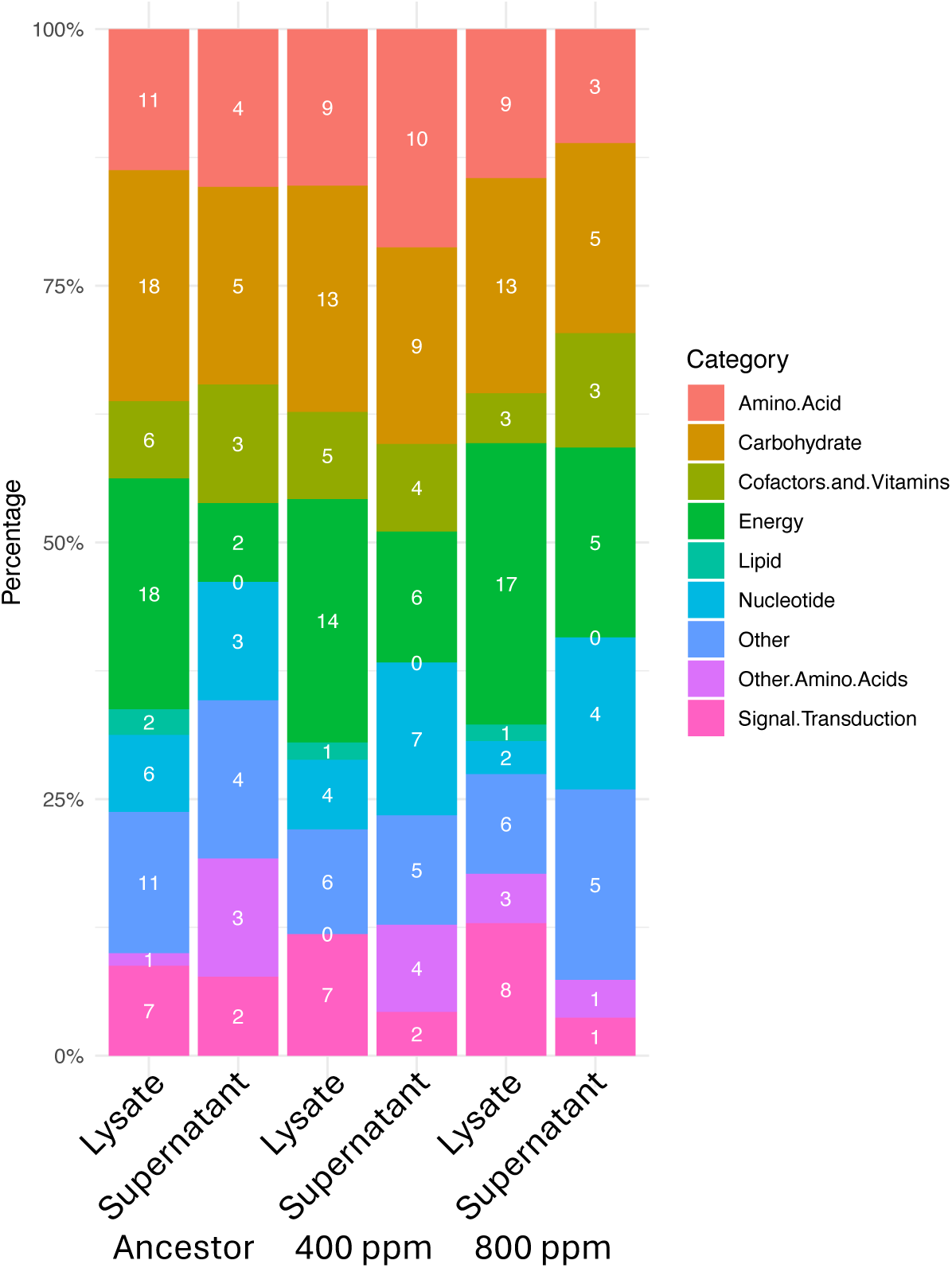
Function of *Alteromonas* proteins. *Alteromonas* strains were isolated from cultures either before (Ancestor) or after 500 generations of evolution at modern or projected future pCO_2_ conditions (400 ppm and 800 ppm, respectively). Proteins were identified by LC-MS from whole cell lysates or from concentrated cell-free supernatants. Discovered proteins were binned into functional categories using BLASTKoala.

**Supplemental Figure 16.**
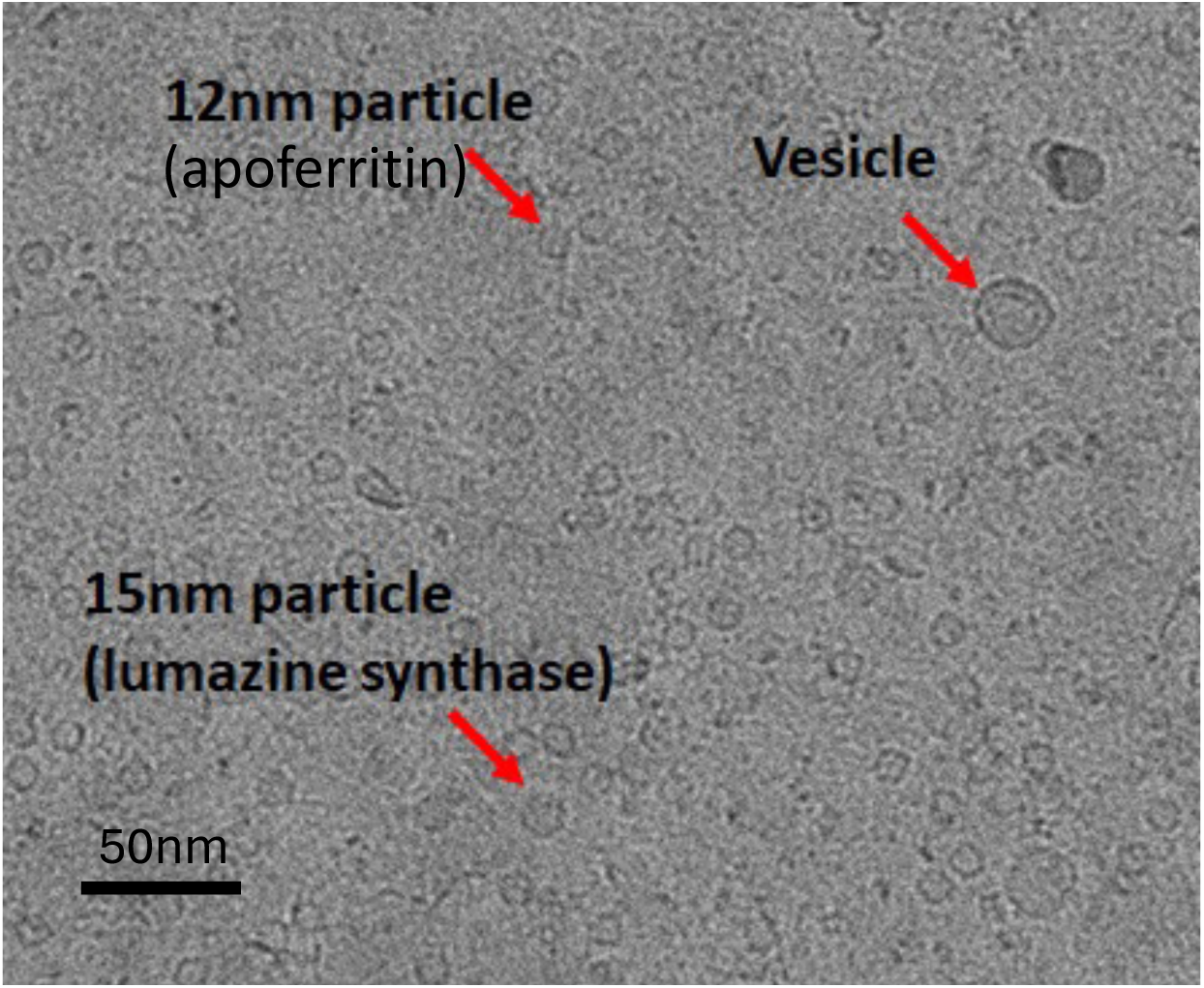
Particles present in *Alteromonas* exudates. In addition to membrane vesicles, cryo-EM also revealed large proteins and protein complexes in the extracellular milieu.

**Supplemental Figure 17.**
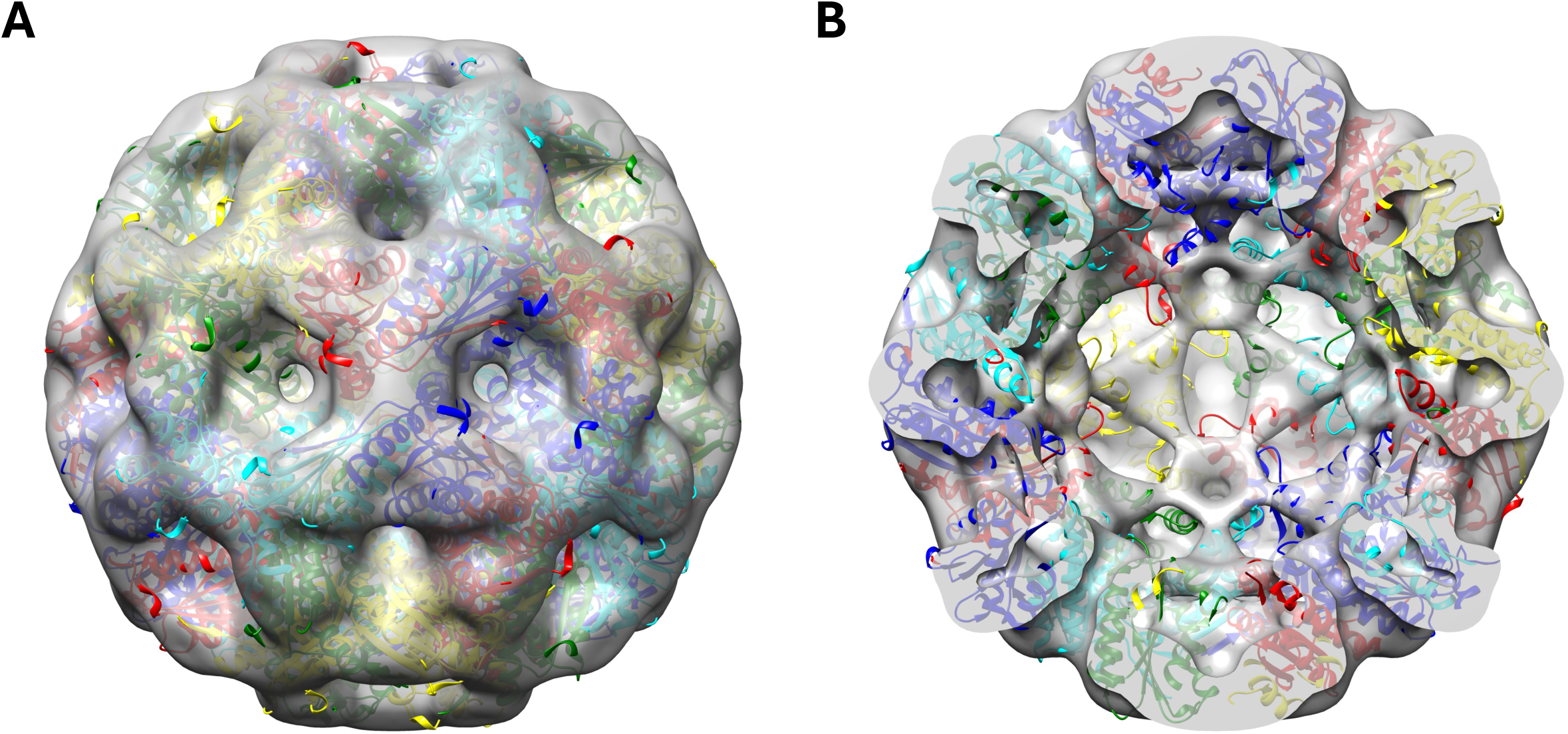
Reconstruction of lumazine synthase from *Alteromonas* exudates. **(A)** The 3D shell generated by Single Particle Reconstruction analysis of the 15nm particle class from cryo EM visualization of *Alteromonas* exudates with the solved crystal structure of *Aquifex aeolicus* lumazine synthase (PDB: 1NQU) superimposed. **(B)** A cross-section of the molecule showing the correspondence of the internal structure with the shape of the protein.

**Supplemental Figure 18.**
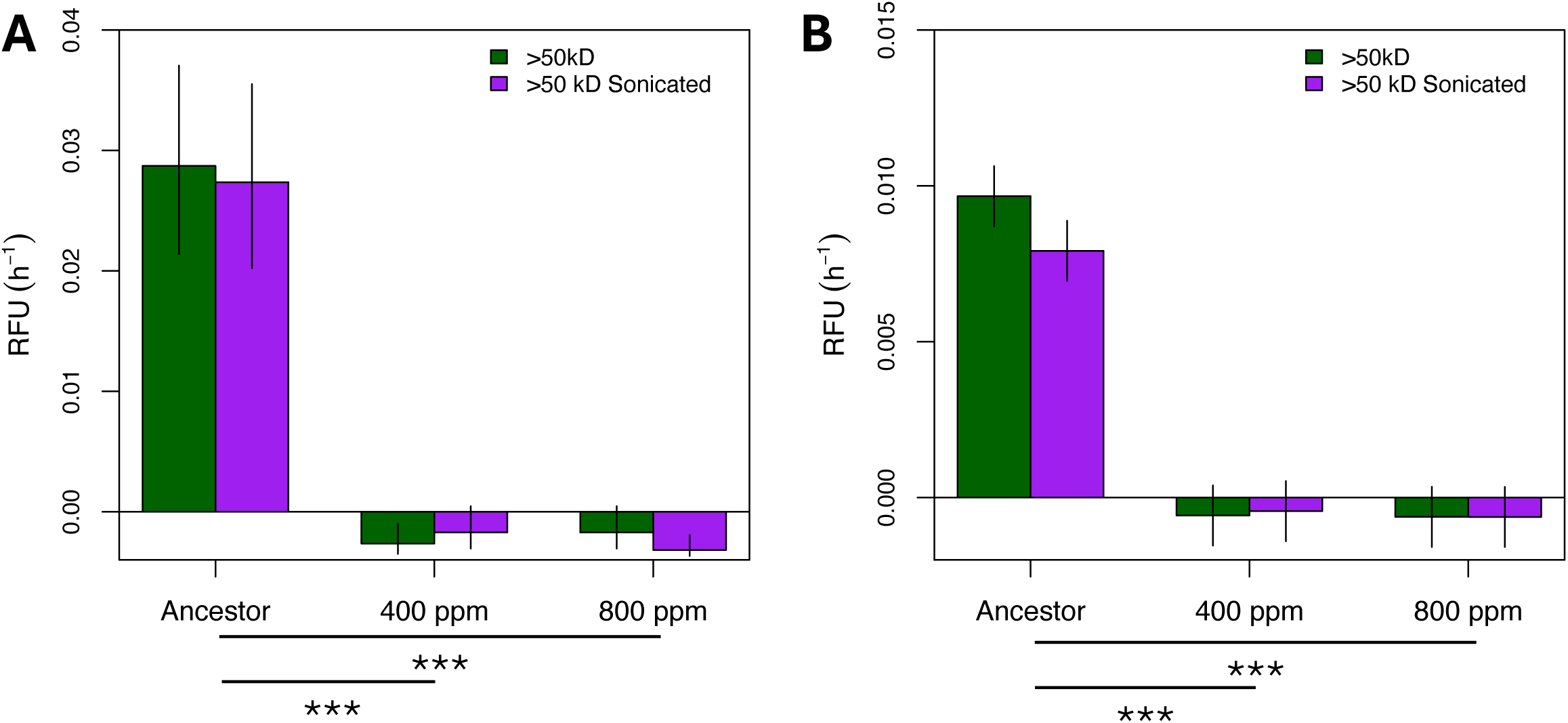
Glycolytic activity in *Alteromonas* exudates. *Alteromonas* strains were isolated from cultures either before (Ancestor) or after 500 generations of evolution at modern or projected future pCO_2_ conditions (400 ppm and 800 ppm, respectively). Cultures were filtered to remove cells and the >50kD fraction was concentrated by tangential flow filtration. Enzyme activities for **(A)** α-glucosidase and **(B)** NAGase were assessed both before and after a sonication treatment to disrupt vesicles. Asterisks below bars indicate between-strain tests using the average of all treatments. ***, *P* < 0.001. No β-glucosidase was detected in the >50 kD fraction.

**Supplemental Figure 19.**
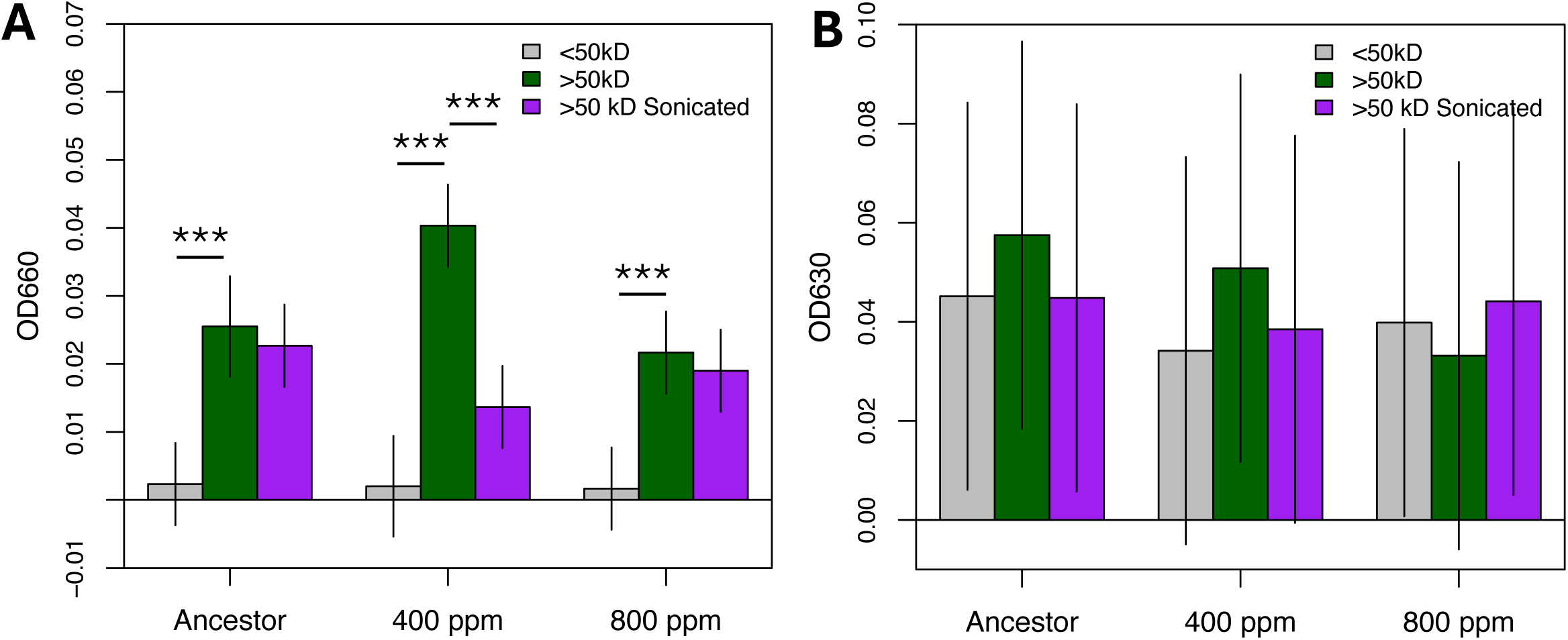
Additional enzyme activity in *Alteromonas* exudates. *Alteromonas* strains were isolated from cultures either before (Ancestor) or after 500 generations of evolution at modern or projected future pCO_2_ conditions (400 ppm and 800 ppm, respectively). Cultures were filtered to remove cells and separated into >50kD and <50kD size fractions by tangential flow filtration. Enzyme activity for **(A)** proteases and **(B)** siderophores were assessed for both fractions as well as for the <50kD fraction after a sonication treatment to disrupt vesicles. All measurements were corrected by subtracting the optical density of a media-only blank; where error bars cross the “0” horizontal, there was no significant difference between the indicated fraction and the blank. Asterisks above bars indicate the results of a within-strain Tukey test. **, *P* < 0.01; ***, *P* < 0.001.

**Supplemental Figure 20.**
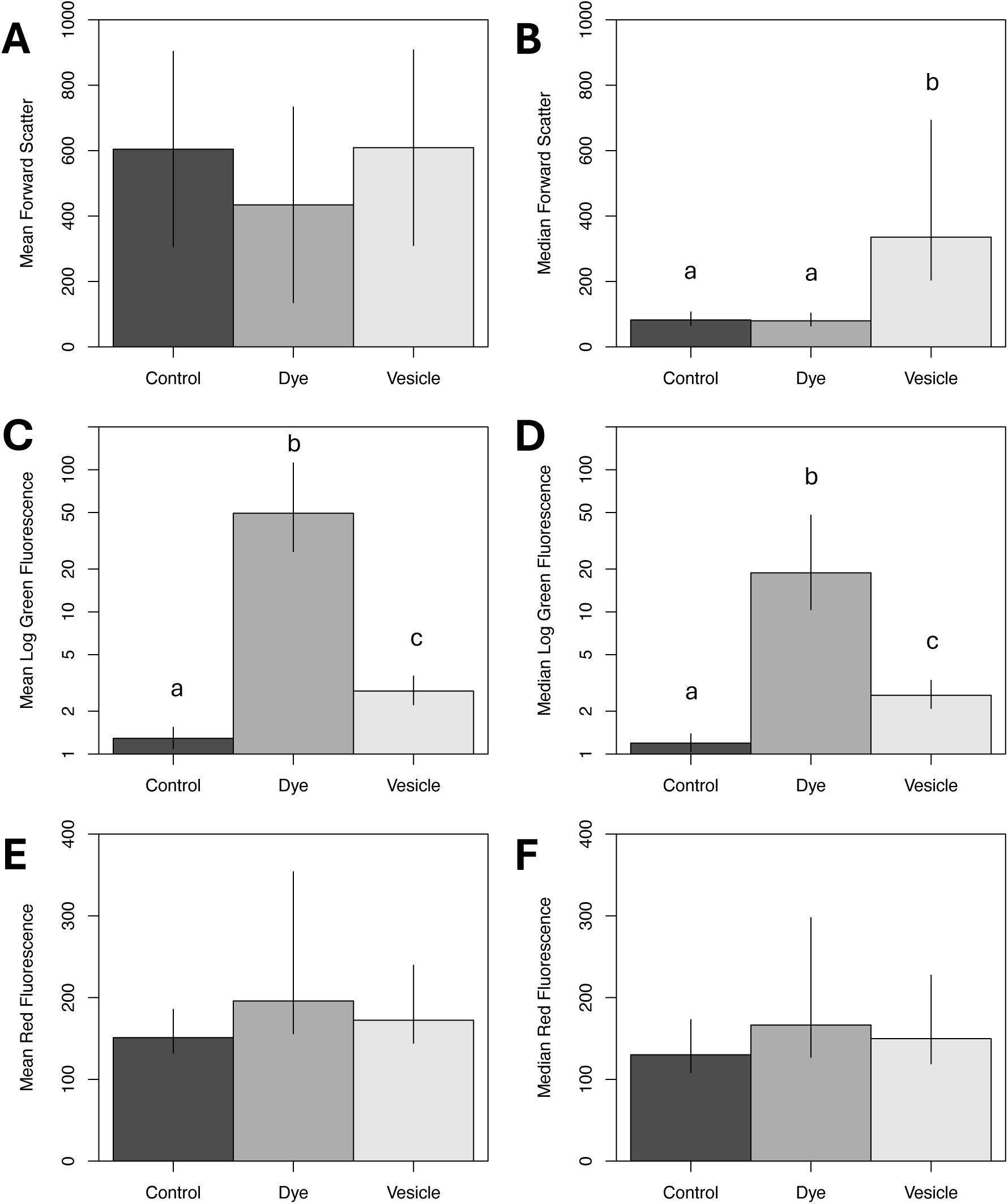
Impact of vesicles on flow cytometry signature of *Prochlorococcus*. *Prochlorococcus* cells were observed by flow cytometry either unmodified (control), after staining with Alexa Fluor 488 5-SDB ester dye (Dye), or after being mixed with similarly stained Alteromonas >50kD concentrates (Vesicle). Error bars indicate the 95% confidence intervals from linear model estimates. Letters above bars indicate groups that are significantly different from each other based on those linear models (p < 0.001); where no letters are present, there were no differences between treatment groups. Values are derived from events in the *Prochlorococcus* gate, i.e., the upper right quadrant shown in Fig. S1A. **(A)** Mean and **(B)** median forward scatter (size); **(C)** Mean and **(D)** median log green (dye) fluorescence; **(E)** Mean and **(F)** median red (chlorophyll) fluorescence

